# Automatic Culture-free Detection of Bacteria from Blood for Rapid Sepsis Diagnostics

**DOI:** 10.1101/2024.05.23.595289

**Authors:** M. Henar Marino Miguélez, Mohammad Osaid, Erik Hallström, Kerem Kaya, Jimmy Larsson, Vinodh Kandavalli, Carolina Wählby, Johan Elf, Wouter van der Wijngaart

## Abstract

Approximately 50 million people suffer from sepsis yearly, and 13 million die from it. For every hour a patient with septic shock is untreated, their survival rate decreases by 8%. Therefore, rapid detection and antibiotic susceptibility profiling of bacterial agents in the blood of sepsis patients are crucial for determining appropriate treatment. The low bacteria concentrations and high abundance of blood cells currently necessitate culture-based diagnostic methods, which can take several days. Here, we introduce a method to isolate bacteria from whole blood with high separation efficiency through smart centrifugation, followed by microfluidic trapping and subsequent detection using deep learning applied to microscopy images.

## 1 Introduction

Sepsis is a severe medical condition characterised by a systemic host inflammation response to infection [1]. Sepsis has an incidence of approximately 50 million cases annually,[2] causing a significant healthcare burden. The mortality of diagnosed patients is around 26% [3]. An estimated 25-30% of sepsis cases involve bloodstream infections (BSI) [4, 5]. The rapid detection and identification of causative organisms in patient blood are important for determining effective antibiotic treatment. The low microbial loads in the bloodstream of patients, as low as one to ten colony-forming units (CFUs) per ml of blood [6], require large sample volumes of blood, which is particularly challenging for pediatric patients [7]. Furthermore, the microorganisms typically need several hours to days of culture before the presence and identification of the pathogens can be confirmed [8, 9]. The latter is problematic, considering the 8% drop in survival rate per hour delayed treatment for patients suffering septic shock [10, 11]. The urgency of the condition, coupled with the latency of the aforementioned diagnosis methods, means that combination therapy with broad-range antibiotics is often prescribed as first-line therapy immediately after the blood drawing [12]. This praxis results in suboptimal treatment [13], contributes to the increase of antibiotic resistance [14], and has also been shown to increase liver toxicity compared to targeted antibiotic monotherapy [15].

The state-of-the-art method for detecting bacteria from sepsis patients is by culturing the blood [16, 17], followed by bacterial identification with genotypic (e.g., polymerase chain reaction), phenotypic (e.g., subcultures), or mass spectrometry methods (e.g., matrix-assisted laser desorption ionisation time-of-flight mass spectrometry (MALDI-TOF)) [18]. While genotypic methods can identify bacteria at low concentrations without culturing the blood [18], they do not give susceptibility information [19], limiting clinical impact. Phenotypic and proteomic approaches typically require a higher bacterial load, necessitating culture and therefore long processing time.

Single-cell phenotypic methods, which specifically bypass the requirement for blood culture, offer a quicker alternative but necessitate the initial removal of blood cells[20–24]. Isolation of bacteria from whole blood has been accomplished with inertial [25–27] and elastoinertial microfluidics [28], sedimentation velocity-based separation [29–31], filtration [29, 32], chemical capture [33, 34], magnetic bead-based separation [35, 36], dielectrophoresis [37], or acoustic separation [38]. However, most of these isolation methods suffer limited throughput and low blood cell rejection rates, or have only been demonstrated for bacteria concentrations of 1000 CFU/ml or above, thus necessitating time-consuming bacterial preculture.

This study aims to develop a rapid, high-throughput assay for isolating bacteria from whole blood with high efficiency and minimal dilution and its concatenation with microfluidic trapping and deep learning-based detection using microscopy images, all within a few hours.

## Results

### Overall work-flow

Our assay concatenates five steps to isolate and detect bacteria from blood: smart centrifugation, selective blood cell lysis, volume reduction, microfluidic trapping combined with miscroscopy imaging, and deep-learning based detection of bacterial cells (Figure 2).

**Fig. 1:**
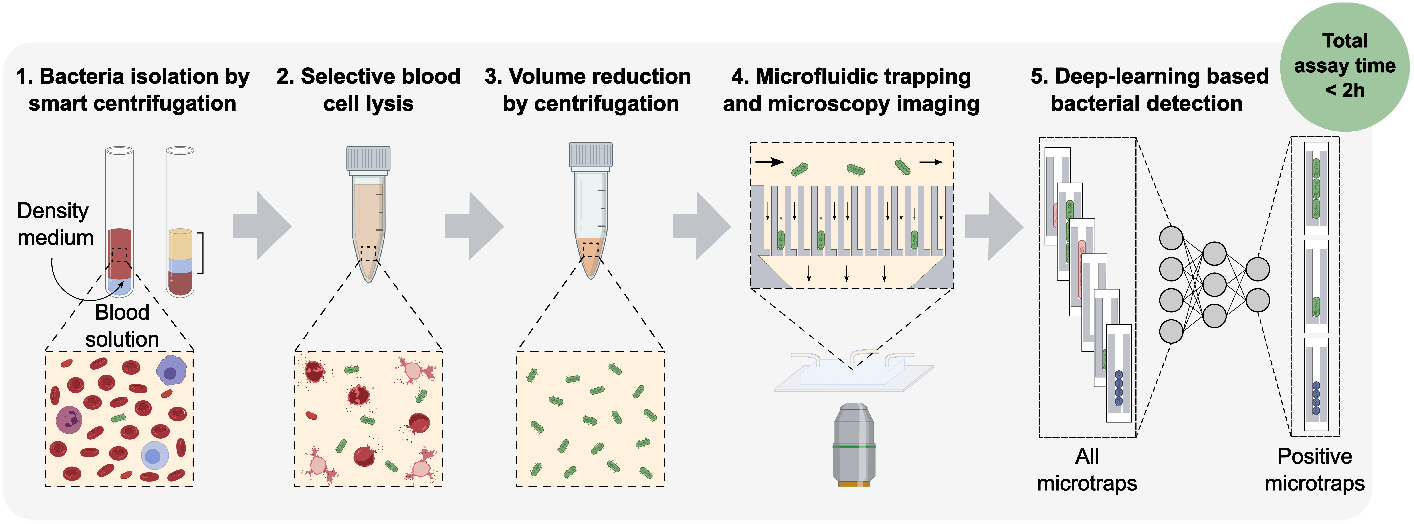
Graphical abstract.

**Fig. 2:**
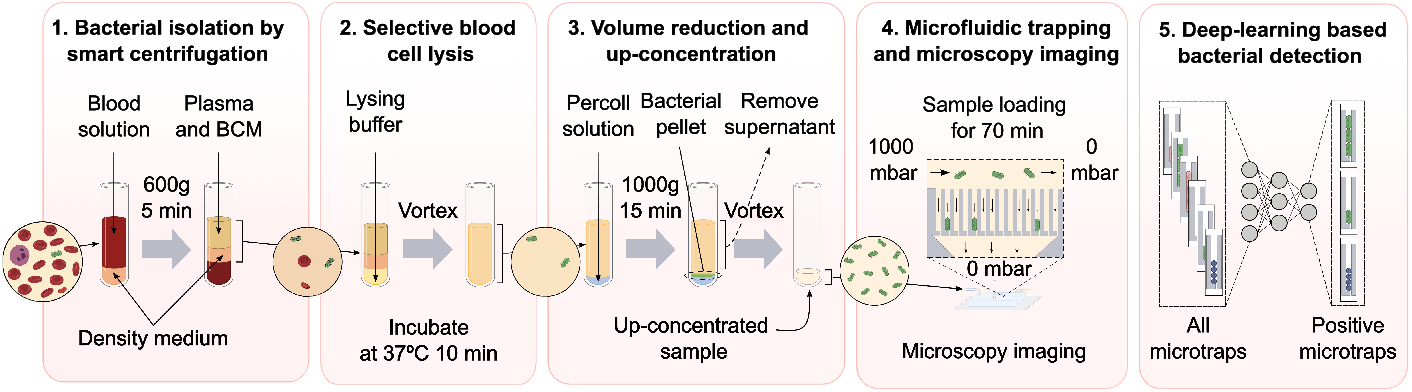
Workflow for bacterial detection from blood samples in five assay steps: 1) isolation using smart centrifugation; 2) selective blood cell lysis; 3) volume reduction; 4) microfluidic trapping and microscopy imaging; and 5) deep-learning based bacterial detection. BCM is blood culture medium.

To allow for repeatability during technical development, this study was conducted with healthy human EDTA donor blood, where we ensured all the blood parameters were within the normal count. The blood was spiked with bacteria at concentrations, *C*, in the range 4-4000 CFU/ml. To facilitate development and quantification, we used a fluorescent lab strain of *E. coli* in the stationary phase (after overnight culture). The fluorescence facilitated the easy distinction between the *E. coli* test strain and potential contaminating bacteria, none of which were observed. Method evaluation was performed with clinical isolates of *K. pneumoniae, E. faecalis*, and *S. aureus* in their exponential growth phase.

Details about materials and methods are provided under “Methods” and all measurement values are tabled in Supplementary information.

### Smart centrifugation

The first assay step, which we call smart centrifugation, removes most of the blood cells while recovering most of the bacteria in the supernatant. This step is essential to avoid downstream clogging of the microfluidic device. During blood centrifugation, bacteria are enriched into the supernatant (see SI for a detailed description of bacterial cell trajectories during blood sedimentation). However, when centrifuging pure whole blood, a large fraction of the bacteria become trapped in the remaining plasma inside the blood cell sediment. Our strategy to avoid such bacterial loss is to layer the sample on top of a density medium with a higher density and with a volume sufficient to replace all plasma in the sediment. We adjusted the volumes and densities of both the sample and density medium for optimal bacterial isolation efficiency with minimal dilution.To tune the densities of the blood sample and density medium, we used blood culture medium (BCM) to support bacterial growth throughout the assay.

The density medium was adapted for an as high as possible density, to increase sedimentation differentiation, with a limit determined by the lowest particle density, to enable the sedimentation of all blood cells into the density medium. The densities of RBCs, WBCs and platelets are in the ranges (1.086 - 1.122 g/ml), (1.057 - 1.092 g/ml) and (1.072 - 1.077 g/ml) [39], respectively; as density medium, we chose a 2:1 volumetric mixture of Lymphoprep (STEMCELL Technologies, Canada) and BCM, which has a density of 1.051 g/ml. The volume of the density medium was experimentally tuned to be minimal but sufficient to replace all plasma remaining between the sedimented cells. Diluting the blood sample with 25% BCM provided minimal reduction of the bacterial concentration while ensuring a sample density below that of the density medium. Centrifuging time and force were experimentally tuned for optimal enrichment of *E. coli* in the supernatant while removing at least 99.8 % of the RBCs. Optimal conditions involved layering 3 ml of BCM-diluted spiked blood on top of 1 ml density medium and centrifuging for 5 min at 600g in a hanging bucket centrifuge. The relative movements of the RBCs, bacteria and liquid in such a system are described with a linear model in Supplementary Information and illustrated in Figure 3A. After centrifugation, approximately 2.5 ml of clear supernatant containing most bacteria (Supplementary Figure 4) could be removed for further processing.

**Fig. 3:**
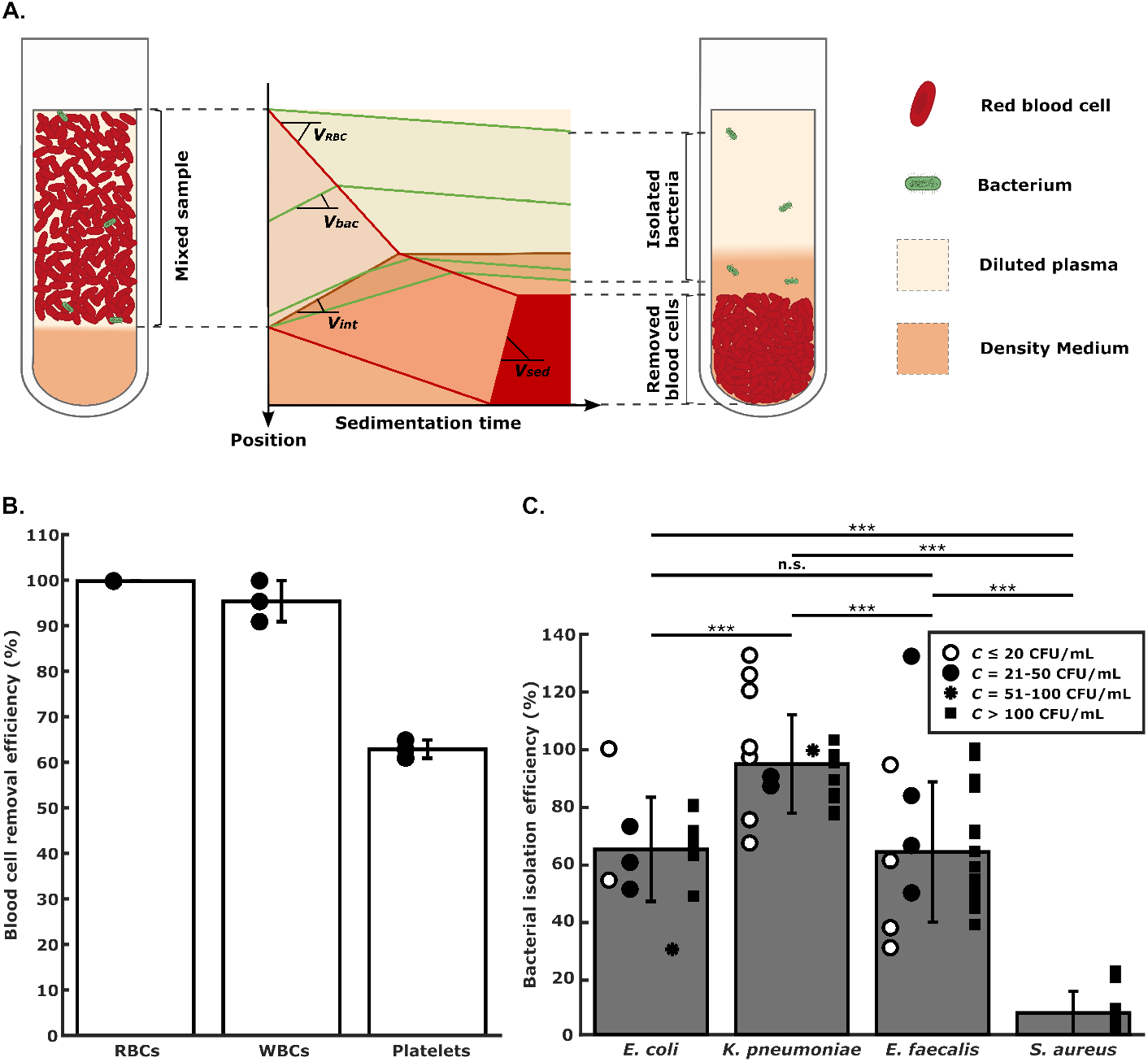
Bacterial isolation from blood by smart centrifugation. **A. Illustration of liquid and cell movement**. The left and right tubes illustrate the positions of sample liquid, density medium, red blood cells and bacteria before and after smart centrifugation. The middle graph qualitatively illustrates the trajectories (solid lines) of bacteria (green) and red blood cells (red) during centrifugation, from a mixed state (left brackets) to a separated state (right brackets). The slopes of the lines are the particle sedimentation speeds, *v*_*RBC*_ and *v*_*bac*_, and liquid interface, *v*_*int*_, and sedimentation interface velocity, *v*_*sed*_, derived in SI. **B. Blood cell removal efficiency**, meaning the fraction of blood cells removed from the supernatant after centrifugation relative to the initial number of blood cells in the sample. **C. Bacterial isolation efficiency**, meaning the number of colony-forming units in the supernatant after centrifugation relative to the initial number of colony-forming units in the spiked sample. Bar heights are mean; error bars are sd; n.s. and *** indicate significance levels p *>* 0.05 and p ≤ 0.001, respectively; the bacterial concentration *C* refers to the CFU /ml in the blood sample.

Blood cell counting of the supernatant showed the removal of 99.82 ± 0.04% of the RBCs, 95 ± 4% of the WBCs, and 63 ± 2% of the platelets (mean ± sd, n=3; Figure 3B). Agar plate culturing of the supernatant revealed the isolation of 65 ± 16% of *E. coli*, 95 ± 17% of *K. pneumoniae*, 64 ± 24% of *E*. ≤ ± ± *faecalis*, or 8 ± 7% of *S. aureus* (mean ± sd, n=10-26; Figure 3C). Recovery exceeding 100% can be attributed to the low plate count numbers, leading to considerable variations [40].

### Selective blood cell lysis

The second assay step removes the remaining blood cells in the sample using a selective lysis mixture of sodium cholate hydrate and saponin [41]. Approximately 2.5 ml supernatant from the smart centrifugation was mixed with 1 ml of the selective lysing solution and kept in a shaking incubator at 37^*°*^C for 10 min, completely lysing remaining RBCs, WBCs and platelets while not affecting bacterial viability.

### Volume reduction

The third assay step enriches the sample and removes the excess lysing buffer in a second centrifugation step. The sample-lysate mixture was layered on top of 0.3 ml high-density liquid, consisting of a 1:2 volumetric mixture of percoll and BCM, and centrifuged for 13 min at 1000g to sediment the bacterial cells on the liquid interface. The supernatant was removed to withhold approximately 0.5 ml liquid containing sedimented bacterial cells, as well as some blood lysate. Sedimenting into a higher-density liquid provided a smoother liquid flow in the microfluidic chip during downstream processing (Supplementary Figure 5), while not significantly affecting the bacterial isolation efficiency (p-value *>* 0.05) (Supplementary Figure 6)

### Bacterial microfluidic trapping

In the last step, we processed the resuspended sample through a microfluidic device with bacterial traps for 70 min while imaging by microscopy. We used the chip design introduced by Baltekin et al. [23] for cross-flow filtration of the bacteria into individual filter traps. The chips contained in total 8000 microtraps of length 50 μm, height 1.25 μm and width 1.25 μm with a restriction of opening 300 nm at their end. Each microtrap can capture bacterial cells and function as a culture chamber. The microtraps were monitored with fluorescence microscopy to detect *E. coli*; alternatively phase contrast microscopy to detect the clinical isolates of *K. pneumoniae, E. faecalis* and *S. aureus*. The 0.5 ml bacterial sample was loaded into the inlet at 1 bar pressure, keeping all outlets at atmospheric pressure.

For *E. coli* -spiked sample, approximately 30% of the sample flowed through the traps with a typical filtrate flow rate of 1-2 μl/min (Supplementary Figure 5B and Supplementary Figure 7A). The trapping efficiency, meaning the number of positive traps, *N*, relative to the number of CFUs after smart centrifugation, was 29 ± 6% (mean ± sd, n=5; Supplementary Figure 7B).

### Microfluidic trapping efficiency

We evaluated the entire assay starting from blood spiked with *E. coli, K. pneumonia*, and *E. faecalis*, respectively. All experiments resulted in the successful trapping and subsequent detection of bacteria. The lowest bacterial concentrations, *C*, tested and detected were 9, 7 and 32 CFU/ml, respectively (Figure 4A). The least mean square linear curve fit, *N* = *η* · *C*, to the data allows estimating the overall assay sensitivity, *η*. We can infer that the limits of detection, which are the bacterial concentrations for which we expect *N* =1 positive trap, *C*_*N*=1_ = 1*/η*, are in the range 1-10 CFU/ml. After trapping, the bacterial cells divide inside the microchannels, while the sample is still being loaded (Figure 4B).

**Fig. 4:**
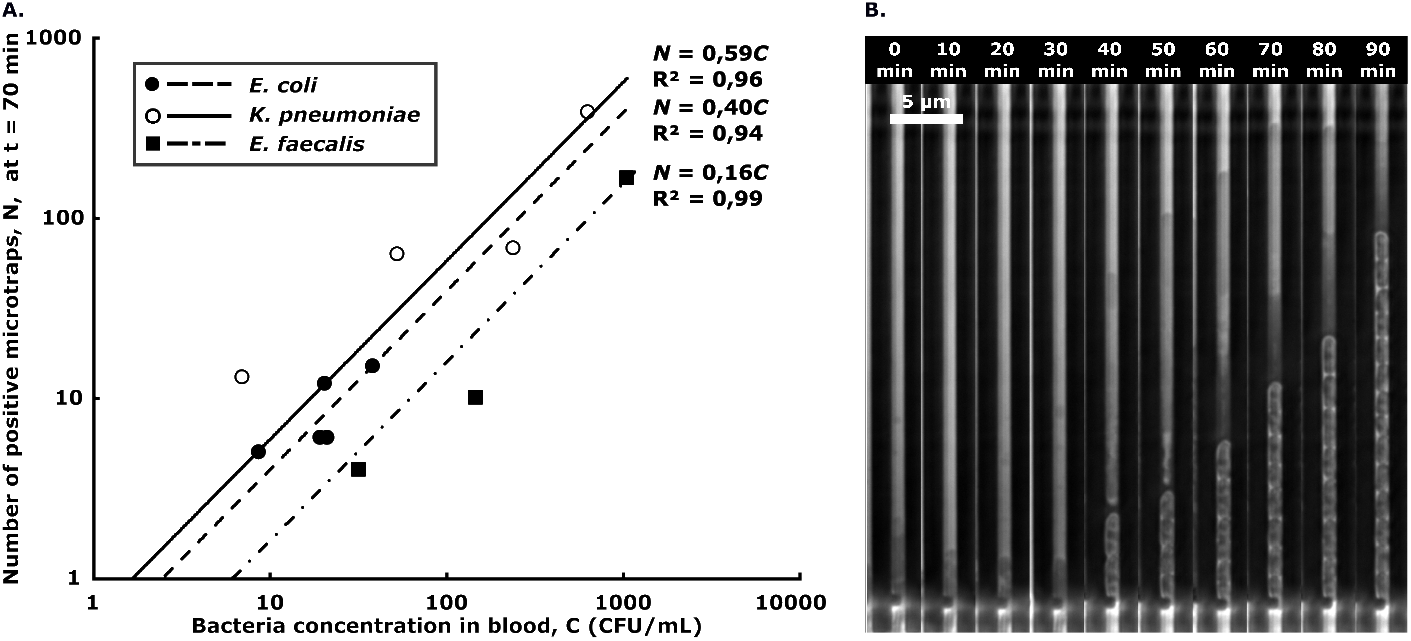
Bacterial microtrapping. **A. Overall assay performance**, where dots represent the number of positive microtraps detected, *N*, for various bacterial concentrations in the 2.25 ml blood sample, *C*, and lines depict the least square fitting linear calibration curves *N* = *η* · *C*. **B. Timelapse images of bacterial capture and growth in a single microtrap**, showing the capture of one bacterium of *K. pneumoniae* 40 min after sample addition, followed by bacterial cell division.

Due to the poor isolation during smart centrifugation, we could not detect *S. aureus* (n=2).

### Deep learning based detection of bacteria

We trained deep learning models to automatically detect the presence of bacteria in 8-frame (70 minutes) 1300×40 pixel (height, width) time-lapses of microfluidic traps, posing the detection as a video classification problem. The dataset was collected through phase-contrast microscopy, capturing bacterial growth in the microfluidic device and then using a pre-processing pipeline to find and crop out individual traps. Each time-lapse was manually labeled, tagging the presence of bacteria in each frame. The training set totaled 57,608 time-lapses (24,220 positives and 33,388 empty at 70 minutes SI Table 6), and the test set 66,869 time-lapses (721 positives and 66,148 empty at 70 minutes SI Table 5), of data from eight experiments each (*E. coli, E. faecalis, K. pneumoniae*). A trap has a positive label if it contains or previously contained any frames with bacteria until the current evaluation time point. The number of positive instances changes over time as cells get trapped in previously empty traps, shown in SI Table 4.

First, time-lapse frames were concatenated horizontally to a time-lapse image, allowing us to use image classification networks for the video detection task. We trained and compared three similarly sized models: ResNet 18[42] from 2016 (11.2M parameters, 8.4 GFLOPs), EfficientNet B2[43] from 2021 (7.7M parameters 3.2 GFLOPs), and the more modern foundation model DinoV2 Small Patch 14[44] from 2023 (21.8M parameters 24.8 GFLOPs). DinoV2 is a standard vision transformer, ViT[45], trained using self-supervision on a large dataset with the DinoV2 methodology; for simplicity, we hereafter refer to it as DinoV2. Then, we employed the Video ResNet R(2+1)D[46] from 2018 (31.3M parameters 47.5 GFLOPs), processing the unaltered timelapse video directly.. The evaluation was conducted by progressively increasing the number of frames in the test time-lapses, measuring the performance over time (0-70 minutes), SI Fig 9A. Also, we trained and tested the models using spatially subsampled image data, simulating a potential clinical setting with a lower-resolution microscope, SI Fig 9B. Each network and subsampling step was retrained 30 times with a specific random seed for reproducibility and better statistics. The evaluation metrics used were precision, defined as: 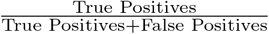, recall (sensitivity), defined as: 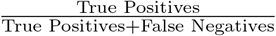, and the F1-score, which is defined as the harmonic mean of precision and recall: *F*_1_ = 2 · 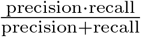. Additionally, ROC (Receiver Operating Characteristic) curves and AUC (Area Under the Curve) scores were calculated. At 70 minutes, the mean and standard deviation F1 scores were 85.5 ± 2.8% for Video ResNet R(2+1)D, 87.3 ± 2.2% for ResNet 18, 90.1 ± 2.3% for EfficientNet B2 and 93.1 ± 1.6% for DinoV2 respectively (full resolution). Fig. 5 shows the confusion matrix, subsampling, and time evaluation heatmap for DinoV2 (the best-performing model), along with the time evaluation classification metrics for all models at full resolution. Inference latency for each model are outlined in SI section “Inference times and computational complexity”. Some cases of obvious incorrect manual labeling of the test data were corrected, for details, see SI section “Data cleaning and label adjustments.”

**Fig. 5:**
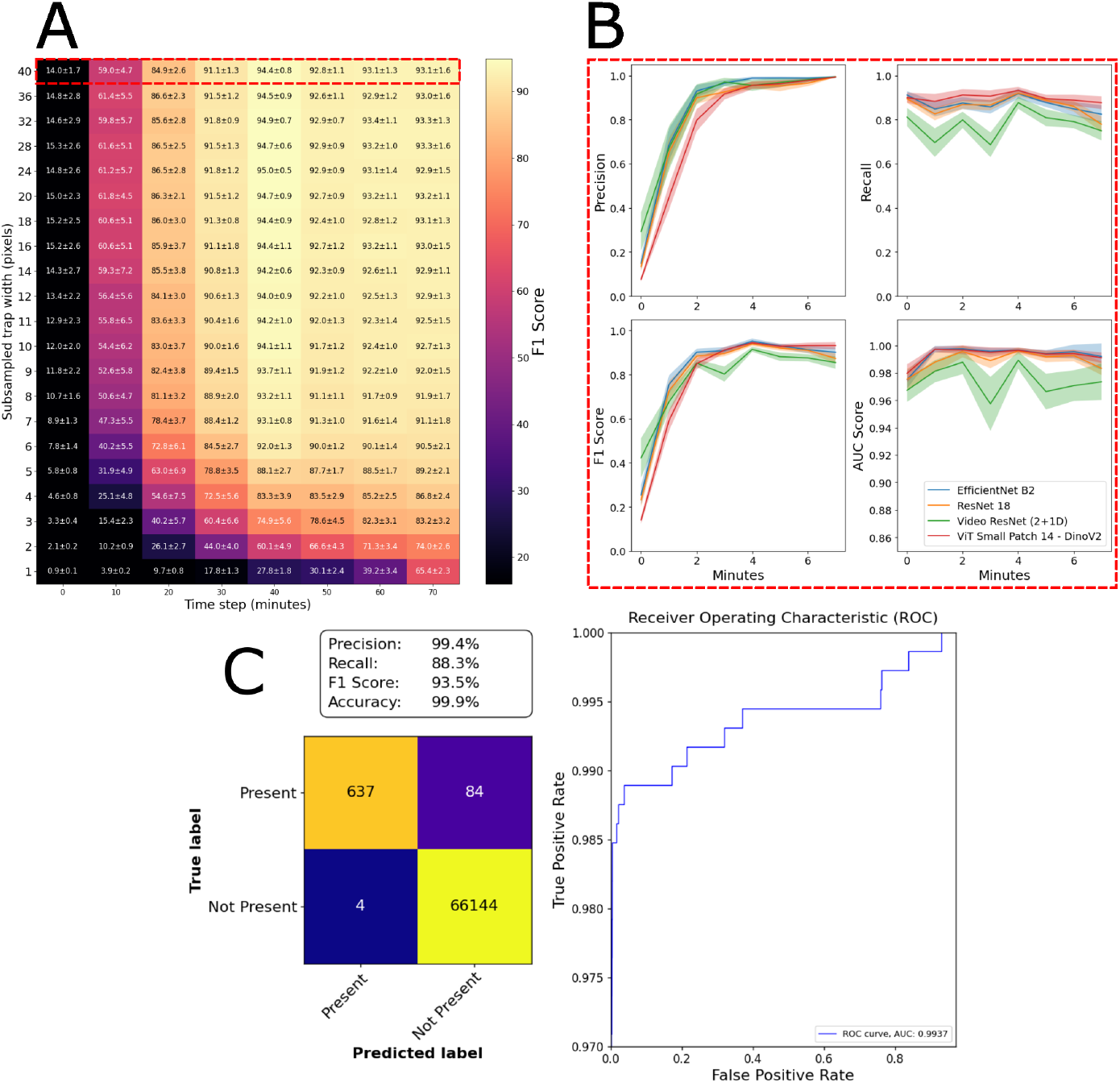
Deep learning model performance. A: Evaluating networks using subsampled data, testing on an increasing number of frames (each corresponding to 10 minutes). The heatmap shows the F1-Score of DinoV2. B: Precision, recall, AUC (Area under the curve), and F1-Score of all models over time at full resolution, corresponding to the red dashed rectangle in part A. Lines show mean, and shaded areas show standard deviations among the 30 retrainings. C: Confusion matrix, performance metrics, and ROC (Receiver Operating Characteristics) curve of the DinoV2 model with the median AUC score among the 30 retrainings, at full resolution, evaluated at the final timestep (70 minutes).

We also performed experiments without considering the time-lapses, performing inference on each frame separately, outlined in SI “Posing the problem as single-frame image classification.” This approach is also feasible and demonstrates good performance, as shown in SI Fig. 13.

## Discussion

We demonstrate the unique isolation and concentration of three sepsis-causing bacterial species from blood, coupled to their automated detection by microscopy in microfluidic traps within 2 h. These pathogens cover approximately 20% of the bacterial species causing BSIs in the European Union [47] and 35% of those in Japan [48]. The estimated detection limit in the range 1-10 CFU/ml is clinically relevant. Compared to previous work, our approach does not require blood culture and allows the detection of bacteria phenotypically at clinically relevant concentrations.

Smart centrifugation is a simple, inexpensive, high-throughput, and easily parallelizable method to isolate bacteria from whole blood rapidly with a high bacterial isolation efficiency and high blood cell rejection. The method’s compatibility with standard lab equipment supports translation to the clinical setting. The bacteria are isolated in a small volume of liquid, similar to the original plasma volume, which facilitates downstream processing.

Bacterial enrichment in the supernatant during centrifugation results from two effects. First, the Stokes terminal velocity of bacteria is a factor *α* ≃ 1/30 times that of blood cells, allowing efficient terminal velocity-based differentiation.[30] Second, the high hematocrit results in downward sedimenting RBCs causing a backflow of the plasma, thereby dragging bacteria upward and further increasing differentiation between the two. Smart centrifugation enhances the separation by extending the sedimentation path of particles through the density medium. For comparison, we experimentally optimised isolation of *E. coli* from spiked whole blood in conventional centrifugation, i.e., without sample dilution or density medium, and achieved 34% ± 7% isolation efficiency in the supernatant (Supplementary Figure 2). Smart centrifugation thus provides a 65% / 34% = 1.9 times improved isolation efficiency for *E. coli* for a comparable blood cell rejection efficiency (Supplementary Table 3).

Even though our method was optimised for *E. coli*, we achieved even higher isolation efficiency of *K. pneumoniae*. Isolation efficiency of *S. aureus*, however, was much lower. Isolation efficiency of *E. coli* and *E. faecalis* were not significantly different. We found no significant differences in bacterial isolation efficiencies between samples spiked with the same strain at different concentrations (Supplementary Figure 1).^1^ We hypothesize that several factors can cause the recovery differences between the species, as well as the large variation in recovery between different measurements with the same strain. These factors include differences in Stokes radius between the bacterial species [30], differences in interaction between the blood cells and the bacteria, differences in viscosity and hematocrit between the blood samples, variations in the immune response of the patient blood to the bacteria (reducing the bacterial count), or variations in bacterial growth phase (compared to *K. pneumonia, E. coli* were spiked in stationary phase) or growth rate (*E. faecalis* have a slower growth rate than *K. pneumonia*). We speculate that the low isolation efficiencyof *S. aureus* is caused by their binding to blood cells or plasma protein, leading to their rapid sedimentation [31]. Indeed, spiking *S. aureus* in a sample liquid with the same density as diluted blood - but without blood cells - resulted in 54 ± 6% (n=6) isolation efficiency by smart centrifugation, compared to 8 ± 7 % (n=10) isolation efficiencyfrom spiked blood (Supplementary Figure 3).

Based on our reading of the literature, smart centrifugation has superior recovery and throughput for *E. coli*, data for which are presented for most tests (Supplementary Table 3).

The concentrated bacterial sample was loaded onto the microfluidic chip at 1000 mbar. At higher pressures, we observed bacterial loss through the 300 nm constrictions of the microtraps. Sedimenting into a higher-density liquid during the prior volume reduction step reduced the compression of bacterial cells and lysate and allowed more homogeneous resuspension, leading to lower on-chip clogging. The trapping of bacterial cells and blood cell debris in the traps increases the fluidic resistance of the filter and decreases the through-filter flow over time. The chips enabled processing 0.5 ml sample within 70 to 90 min, during which 30% of the sample flows through the microtraps (filtrate flow) and the rest flows past the filter directly to the waste outlet (retentate flow). The low filtrate flow rate is thus a major contributor to bacterial loss between isolation and detection.

The deep learning experiments demonstrate the feasibility of using deep neural networks to accurately detect the presence of bacteria in phase-contrast time-lapses. Both transformer-based models and convolutional neural networks achieve similar performance. The task can also be framed as single-frame image classification with good results. However, in a real-world scenario, it would be more advantageous to use the entire time-lapse sequence to detect bacterial growth, especially if using low-resolution microscopy. Inference times for a single time-lapse ranged from 1 to 3 milliseconds on our system, indicating that latency should not pose a significant issue, even on automated microscopy systems with less powerful hardware.

## Methods

### Microfluidic Platform Fabrication

The microfluidic chip design and fabrication were previously reported [23, 49]. A silicon wafer mold was fabricated according to design specifications by the company ConScience AB, Sweden. The wafer was once silanized for 30 min prior to polydimethylsiloxane (PDMS; Sylgard 184, DOW, USA) replication. Subsequently, a 10:1 w/w PDMS:curing agent mixture was poured on the wafer and baked at 80°C overnight. Holes on the PDMS ports 2.0, 2.1, 2.2, 5.1 and 5.2 (see chip design in [23, 49]) were made using a 0.5 mm puncher. Then, the PDMS stamps were cleaned in IPA before covalent bonding to a glass coverslip (No. 1.5, Menzel-Glöaser, Germany) after plasma treatment, followed by heat curing for 1 h at 80°C.

### Microfluidic Flow Control

The microfluidic device was mounted on the microscope and connected to the sample reservoirs with flexible plastic tubing (TYGON, Saint-Gobain, North America). OB1 CONTROLLER (Elveflow, France) flow control units were used to pressurize reservoirs.

The reservoirs connected to ports 2.1, 2.2, 5.1 and 5.2 were first pressurized at 500 mbar for chip priming with water infused with 0.085 g/l of Pluronic F108 (Sigma-Aldrich, USA) and was decreased to 0 mbar. Then the reservoir connected to inlet port 2.0 was pressurized at 500 mbar for priming and at 1000 mbar for sample loading.

### Blood Sample Preparation

Blood in EDTA tubes from healthy donors was purchased from the blood bank (Blodcentralen, Stockholm, Sweden) or Uppsala University Hospital (Akademiska Sjukhuset, Uppsala, Sweden). The blood samples were stored at 4 ^*°*^C, and used for experiments no later than two days after collection. Samples presenting a milky supernatant (*<* 10%), i.e., indicating abnormally high concentrations of triglycerides in the blood, were discarded. Before processing the samples, they were put at room temperature, and blood samples were spiked with respective bacteria concentrations of approximately 10^3^, 10^2^, and 10 CFU/ml. The concentration of the spiking solution was quantified by plate counting (n=3). The spiking volume was always less than 5% of the total blood solution volume.

### Bacterial Strains

We used three clinical isolates that were randomly collected from a clinical microbiology laboratory in Sweden, covering both gram-negative and gram-positive species. As gram-positive representatives, *S. aureus* (DA70300) and *E. feacalis* (DA70208) were used, and as gram-negative, *K. pneumoniae* (DA72206). We also used *E. coli* (EL3041) cells harbouring a plasmid expressing mVenusNB fluorescence proteins.

Bacteria were stored for long term at −80 ^*°*^C in standard glycerol solution and were incubated overnight at 37 ^*°*^C in BD BACTEC Plus Aerobic medium (BD, USA), referred to as Blood Culture Medium (BCM), prior to use, and later diluted to approximately 10^4^, 10^3^, and 10^2^ CFU/ml and used to spike blood with different concentrations and quantified for isolation. For detection experiments with *E. coli* cells harbouring a plasmid expressing mVenusNB fluorescence proteins, the above-mentioned protocol was followed. For detection experiments with clinical isolates, 2 μL of bacteria from an overnight culture were spiked in 2 ml of BD BACTEC Plus Aerobic Medium, and cultured for 2-4 h depending on the species. They were then diluted to approximately 10^4^, 10^3^, and 10^2^ CFU/ml and used to spike blood with different concentrations.

### Smart Centrifugation

Lymphophrep (STEMCELL Technologies, Canada), a density medium with 1.077 g/ml density, was mixed with BD BACTEC Plus Aerobic medium (BD, USA) in a volumetric ratio of 2:1, and 1 ml of the solution, with 1.051 g/ml density, was poured into a 15 ml Falcon centrifuge tube. 3 ml of spiked blood mixed with BD BACTEC Plus Aerobic medium (BD, USA) (3:1 Blood:BD BACTEC Plus Aerobic medium) was gently placed over the density media. The tube was centrifuged at 600g for 5 min in a hanging bucket centrifuge. The supernatant was removed and mixed with the lysing solution.

### Conventional centrifugation for bacteria isolation

To establish a control measurement, we centrifuged 4 ml of whole blood without any dilution and density media. The sample, after being spiked, underwent centrifugation at 500g for 4 min, resulting in the separation of clean plasma at the top. This process effectively settled most of the blood cells.

### Selective Lysing Solution

The selective lysing solution was prepared by mixing sodium chocolate hydrate (Sigma-Aldrich, USA) and saponin (Sigma-Aldrich, USA). Concentrations of 2% (W/V) sodium cholate hydrate and 1% (W/V) saponin were prepared by dissolving the chemicals in BD BACTEC Standard Aerobic medium.

### Volume reduction

Percoll (Sigma-Aldrich, USA), a density medium having a density of 1.125-1.135 g/ml was mixed with BD BACTEC Plus Aerobic medium (BD, USA) in a volumetric ratio of 1:2, and 0.3 ml of the solution, with approximately 1.04 g/ml density, was poured into a 15 ml Falcon centrifuge tube. The supernatant infused with the lysing solution was gently placed over the density medium. The tube was centrifuged at 1000g for 13 min in a fixed rotor centrifuge. All excess liquid above 0.5 ml was removed, without removing the pellet.

### Optical setup

Images of all the microtraps contained in the microfluidic device were acquired using a Nikon Ti, inverted microscope. For imaging experiments containing clinical strains, a CFI plan Apo lambda 100x (1.45 NA, oil) objective was used. Phase contrast images were acquired by using DMK 38UX304 (the imaging source) camera, with an exposure time of 20 ms with an interval of 10 min, for a maximum of 90 min. Cells expressing mVenus fluorescence proteins were captured using a filter cube consisting of a Di02-R488 (Semrock, USA) dichroic mirror, a FF02-482/18 (Semrock, USA) excitation filter, and a FF01-524/24 (Semrock, USA) emission filter and CFI Plan Apo lambda DM Ph2 20x objective. Each fluorescence image was acquired with an exposure time of 500 ms with an interval of 5 min, for a maximum of 90 min. A constant temperature of 37ºC was maintained during the measurements using a temperature-controllable unit (Okolab, Italy). The imaging setup was operated by micromanager 1.4 version software.

### Bacteria Quantitation

All bacterial sample quantitation was performed by plate counting after plating a sample aliquot on agar plates and overnight culture at 37 ^*°*^C in the incubator. The agar plates were prepared by dissolving LB broth with agar (Miller) (Sigma-Aldrich, USA) in deionized water at 40 g/L concentration followed by autoclaving and placing in Petri dishes.

For on-chip bacteria quantitation of the clinical isolates, we employed ImageJ software image analysis of phase-contrast microscopy images. Each microtrap was analyzed manually and determined positive if filled with at least one bacterium. The same analysis pipeline was performed for fluorescence microscopy images of the *E. coli* cells expressing mVenus fluorescence proteins.

### Blood Cell Quantitation

The concentrations of the blood cells in whole blood before dilution and in the supernatant after smart centrifugation were measured using a haematology analyzer (Swelab Alfa Plus, Boule Diagnostics, Sweden).

### Flow characterization

The ports 2.1 and 2.2 were coupled together, and ports 5.1 and 5.2 were coupled together, using a PEEK union T-connector (Thermo Fisher Scientific, USA). Flow measurements were taken from both couplers using a SLI Liquid Flow Sensor (80 μL/min) (Sensirion, Switzerland). Two independent flow measurements were acquired with a sampling time of 1 s, for a maximum time of 90 min, using the Sensor Viewer Software (Sensirion, Switzerland). The data was analysed using MATLAB.

### Image Cropping and stabilizing

An image processing pipeline was further improved from [50] to rotate, find, and crop out 1300×40 pixel trap images in the microfluidic chip, and then stabilizing the corresponding time-lapse described in SI “Preprocessing and cropping”. During deep learning model training and inference the frames were cropped vertically at 336 pixels from the physical stop at the top of each trap and padded to a width of 42 pixels. The reason for padding the traps to 42 pixels width is that we hypnotize DinoV2 would perform better, as it uses patches of 14×14 pixels for the token embeddings, meaning it would fit three tokens on the width of a trap. The rationale for the 336-pixel cropping of the height was to fit the time-lapses in the GPU memory and concerns of overfitting. Additionally, the difficult instances had one or a few cells situated at the top adjacent to the physical stop, thus, processing the whole trap was unnecessary.

### Time-lapse images and subsampling

When employing image classification models (ResNet 18, DinoV2, and EfficientNet B2), the frames of each trap were concatenated horizontally to a 336×336 pixel 2D image showing the time-lapse. Zero-padding was applied on the right side when performing the time evaluation, shown in SI Figure Padding was also applied during training if the time-lapses from the experiment contained less than eight frames, seen in Table 6. For the Video ResNet R(2+1)D, we instead relied on the global averaging pooling [51] to test and train on time-lapses with different numbers of frames. A time-lapse sample had a positive label if it contained any positive frames to the evaluated time-point (also if cells had been previously trapped and subsequently escaped). Lancoz[52] subsampling was used to simulate using lower-resolution microscopy. Before feeding the subsampled images into the model, they were upsampled to their original size using bicubic interpolation. This step was necessary because DinoV2 is a transformer with a fixed patch size, thus, it could not process images with smaller widths than 14 pixels. The single-channel grayscale phase-contrast data was transformed to RGB using the Vidiris color map [53] for visibility and to aid human labeling. The transformed color images were also used when feeding into the networks (we kept the 3-channel input).

### Network training

All models were trained for 20 epochs with batch size 32 and learning rate 10^*−*3^ except DinoV2 (batch 128, learning rate 10^*−*5^, trained for 15 epochs) and Video ResNet R(2+1)D (used learning rate 10^*−*4^). All networks used pretrained weights, employed the standard ADAM optimizer[54], Cross Entropy loss, Cosine learning rate scheduler[55], and 15% label smoothing [56]. The Pytorch deep learning framework was used with Pytorch Image Models (Timm)[57]. During training, images were sampled with replacement for each mini-batch from the two classes, with sampling probabilities inversely proportional to the class overrepresentation in the training set (using a Random Weighted Sampler in Pytorch). The augmentations ShiftScaleRotate, HorizontalFlip, VerticalFlip, RandomBrightnessContrast, and Blur were employed during training from the Albumetations library[58]. Additionally, 1-7 frames (sampled uniformly) were randomly erased during training (random frame erase augmentation). All models were trained on the NvidiaA100 GPU.

For further information, refer to the released software package containing the dataset and software to reproduce the deep learning experiments [59]

### Statistical analysis

Data analyses were performed with R. The data samples were tested for normality using the Shapiro-Wilk test; variance between normal populations was studied using the F-test; and statistically significant differences between groups were studied using a t-test, for samples following a normal distribution, or a Mann Whitney test for samples not following a normal distribution. A two-sided p-value of ≤ 0.05 was considered statistically significant.

## Footnotes

1 With the exception of *E. coli*, where we found a significantly lower isolation efficiency for concentrations from 21 - 100 CFU/ml compared to *>* 100 CFU/ml (p 0.05). However, the statistical power of this finding is low considering the very low number of measurements.

## Notes

### Competing Interest Statement

Yes there is potential Competing Interest.
The authors aim to secure IP protection for at least parts of this manuscript. JE and WW have commercial interests in the diagnostics field including IP but not directly related to isolation of bacteria from blood

### Summary of Updates

This version of the manuscript has been revised to incorporate significant methodological advancements and analysis improvements. We introduced an automatic, culture-free bacterial detection method from whole blood, aiming at rapid sepsis diagnostics. The manuscript now includes detailed integration of a deep learning-based detection step, applied to microscopy images from microfluidic traps, enabling automated identification of bacteria. The manuscript's abstract and workflow have been updated to clearly reflect the integration of deep learning techniques into the previously established isolation and trapping methods. Specifically, the assay now comprises five clearly defined steps: smart centrifugation, selective blood cell lysis, volume reduction, microfluidic trapping combined with microscopy imaging, and deep learning-based bacterial detection. The results section has been expanded to describe in detail the performance evaluation of various deep learning models (ResNet18, EfficientNet B2, DinoV2 Small Patch 14, and Video ResNet R(2+1)D). Performance metrics such as precision, recall, F1-score, ROC curves, and AUC scores are comprehensively analyzed and reported, demonstrating that DinoV2 achieved the highest accuracy. The manuscript now includes additional figures and data supporting the robustness of the AI-driven microscopy approach, including sensitivity analysis under reduced image resolution conditions. Minor textual clarifications and supplementary methodological details have also been added throughout the manuscript to enhance clarity and reproducibility. Updated supplementary information contains comprehensive data tables, detailed descriptions of image preprocessing techniques, model training specifics, and statistical analyses conducted. Overall, the manuscript revisions significantly enhance the technological depth, validation rigor, and translational potential of the described rapid sepsis diagnostic method.

