## Supplementary for "Automatic Culture-free Detection of Bacteria from Blood for Rapid Sepsis Diagnostics"

### Supplementary information

#### Cell and fluid movement during smart centrifugation

The movement of blood cells during centrifugation has been studied extensively.[60] To understand the displacement of bacteria during centrifugation, we here consider a linear model for the simultaneous sedimentation of two types of particles, bacteria and RBCs, through two types of density-layered liquids, diluted plasma and density medium, in which we neglect particle-particle interactions. This manuscript uses the indices *bac*, *RBC*, *pl*, *DM*, *int*, and *sed*, to indicate bacteria, RBCs, plasma, density medium, liquid:liquid interface and sediment, respectively.

The theoretical stokes terminal velocity, $v_{st}$, of particles in blood can be deduced from Stoke's law, which describes the unhindered sedimentation velocity of a single particle in a standing fluid. The stokes velocity depends linearly on the density difference between the particle and the medium, $\Delta\rho$, quadratically on the particle size, and inversely linear on the viscosity, $\mu$. The stokes velocities of a given particle in the two liquids thus relate as

$$\frac{v_{st,DM}}{v_{st,pl}} = \frac{\mu_{pl}.\Delta\rho_{DM}}{\mu_{DM}.\Delta\rho_{pl}}. \tag{1}$$

Because the densities of bacteria and RBCs are in the same range, the ratio of their stokes velocities can be considered a constant, $\alpha$, independent from the liquid medium:

$$\alpha = \frac{v_{st,bac}}{v_{st,RBC}}. \tag{2}$$

$\alpha$ has been estimated to be typically 1/30 [30].

In regions with a large local volume fraction of RBCs, $h$, the RBC sedimentation causes a significant upward backflow of liquid medium. Considering RBCs incompressible, the local backward liquid velocity, $v_l$, can be deduced from mass conservation, i.e., the downward flux of RBCs must equal the backward flux of liquid:

$$h.v_{RBC} + (1-h).v_l = 0 \Longleftrightarrow v_l = -\frac{h}{1-h}.v_{RBC}. \tag{3}$$

(Throughout this manuscript, we use a negative sign for velocities to indicate an upward movement direction opposite to that of downward sedimenting RBCs.) Where the sedimenting RBCs cross the liquid interface between the sample and density medium, mass conservation demands that the local liquid velocities across the interface, and the velocity with which the interface itself moves backward, are all equal. This equality implies a relation between the difference in local RBC fractions, $h_{pl}$ and $h_{DM}$, across the liquid interface:

$$v_l = v_{int} = v_{pl} = v_{DM} = -\frac{h_{pl}}{1-h_{pl}}.v_{RBC,pl} = -\frac{h_{DM}}{1-h_{DM}}.v_{RBC,DM}. \tag{4}$$

The effective sedimentation velocity of the particles equals the sum of the local liquid velocity and their theoretical Stokes velocity. For RBCs, the resulting effective sedimentation velocity can be expressed as

$$v_{RBC} = v_l + v_{st,RBC} = -\frac{h}{1-h}.v_{RBC} + v_{st,RBC} \Longleftrightarrow v_{RBC} = (1-h).v_{st,RBC}, \tag{5}$$

i.e., $v_{RBC,pl} = (1 - h_{pl}).v_{st,pl,RBC}$ and $v_{RBC,DM} = (1 - h_{DM}).v_{st,DM,RBC}$.

The values for $v_{st,RBC}$ can be derived relative to those of blood using equation 1. The value of $h_{pl}$ can be estimated from the hematocrit value while accounting for the 25 % blood sample dilution with BCM. The value of $h_{DM}$ can then be derived from $h_{pl}$ using equation 4.

During sedimentation, bacteria can assume any of four velocities, depending on their location in either the plasma or density medium, and on the local RBC fraction being zero or not.

$$v_{bac} = v_l + v_{st,bac} = -\frac{h}{1-h}.v_{RBC} + \alpha.v_{st,RBC} = -(h-\alpha).v_{st,RBC}. \tag{6}$$

In the supernatant and in density medium depleted from RBCs, $h = 0$ and $v_{pl} = 0$ and bacteria reach their stokes terminal velocity $v_{bac} = v_{st,pl,bac}$.

Importantly, and perhaps counterintuitively, bacteria move upwards where bacterial fractions $h > \alpha$.

The speed at which the sedimentation layer grows depends on the liquid fraction, $\beta$, remaining between the RBCs in the sediment:

$$v_{sed} = v_l.(1+\beta). \tag{7}$$

The values of $v_{int}$, $v_{bac}$, $v_{RBC}$ and $v_{sed}$ are visualised as the slopes of the position-versus-time trajectories indicated in Figure 3A.

#### Statistical Analysis of Smart Centrifugation of Samples with Different Bacterial Concentration

We have studied the variation in isolation efficiency of bacteria by smart centrifugation between the different concentration ranges < 20, 20-100 and >100 CFU/ml for three bacterial species (Supplementary Figure 1). For *K. Pneumoniae* and *E. Faecalis* we found no significant difference. For *E. coli* we found a significant difference only in between the concentration ranges < 20 CFU/ml and 20-100 CFU/ml.

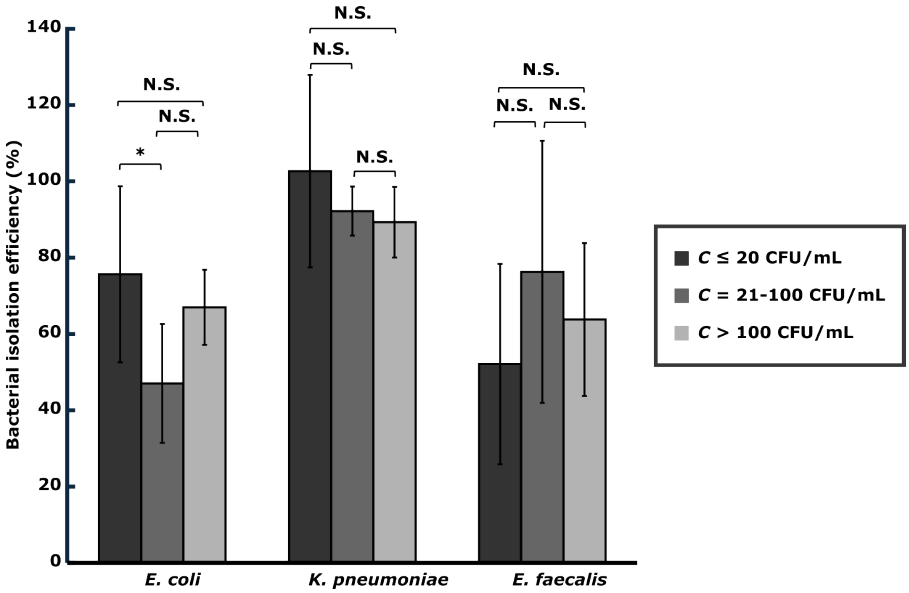

**Supplementary Figure 1**: **Isolation efficiency of bacteria from blood by smart centrifugation in different concentration ranges.** n.s. and * indicate significance levels $p > 0.05$ and $p \leq 0.05$, respectively.

#### Isolation of bacteria by conventional centrifugation of undiluted blood without density medium

The bacterial isolation efficiency for (*E. coli*) has also been studied for conventional centrifugation, i.e., centrifugation without sample dilution or density medium, of whole blood. 4 ml of whole blood was centrifuged at 500g for 4 min. 4 min was sufficient for most of the blood cells to settle. We recovered 34 ± 7 % (mean ± sd; n=3) of the bacteria in the supernatant (Supplementary Figure 2), removing 99.957 ± 0.007% of the RBCs, 98.9 ± 0.6 % of the WBCs, and 65 ± 2 % of the platelets (mean ± sd; n=3) (Supplementary Table 1).

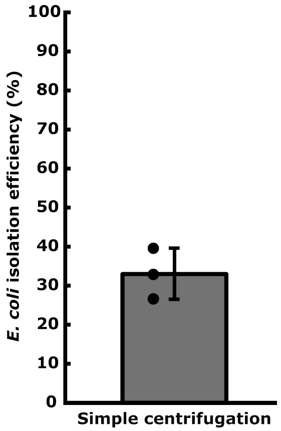

**Supplementary Figure 2**: **Bacterial isolation efficiency in conventional centrifugation of blood.**

**Supplementary Table 1**: Data points for % rejection of blood cells in conventional centrifugation.

| Cell type | % rejection (E1) | % rejection (E2) | % rejection (E3) | % Average rejection | SD |
|---|---|---|---|---|---|
| RBC | 99.94 | 99.96 | 99.96 | 99.953 | 0.012 |
| WBC | 99.19 | 98.13 | 99.26 | 98.86 | 0.63 |
| Platelets | 62.76 | 65.86 | 67.22 | 65.3 | 2.3 |

### Blood cell rejection in smart centrifugation

**Supplementary Table 2**: Data points for % rejection of blood cells in smart centrifugation.

| Cell type | % rejection (E1) | % rejection (E2) | % rejection (E3) | % Average rejection | SD |
|---|---|---|---|---|---|
| RBC | 99.87 | 99.8 | 99.79 | 99.820 | 0.044 |
| WBC | 90.9 | 95.27 | 99.86 | 95.3 | 4.5 |
| Platelets | 63.27 | 64.68 | 60.71 | 62.9 | 2.0 |

#### Low isolation efficiency of *S. aureus*

We investigated whether the poor isolation efficiency of *S. aureus* is caused by their interaction with the blood cells or plasma protein, as reported by many, or whether the poor performance is intrinsic to smart centrifugation. We replaced the spiked blood in smart centrifugation with a control liquid having the same density and volume as the diluted sample, and performed smart centrifugation by placing it over 1 ml density media. After 5 min of centrifugation, the top 2.5 ml of liquid was removed and plated on an agar plate for bacteria quantification. We recovered approximately 55% of the bacterial cells, whereas the isolation efficiency from blood was less than 10% (Supplementary Figure 3). This difference indicates that the interaction between *S. aureus* and blood cells is stronger than the fluid drag, transporting the bacteria into the sediment .

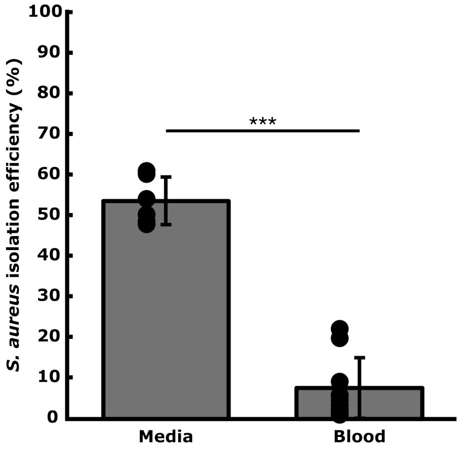

**Supplementary Figure 3: Isolation efficiency of *S. aureus* from blood with and without argatroban and the same density control liquid in smart centrifugation.** *** indicates significance levels p $\leq$ 0.001.

#### Distribution of bacteria in the supernatant and pellet after smart centrifugation

We investigated the distribution of *E. coli* in the pellet and the supernatant after smart centrifugation. After the centrifugation step, the supernatant was removed and the pellet was resuspended in 2 ml of BCM to make plate counting more accurate as a very high number of interfering blood cells can decrease the accuracy of plate counting. We recovered almost all (> 90%) of the spiked bacteria. 30% of the bacteria were found in the pellet.

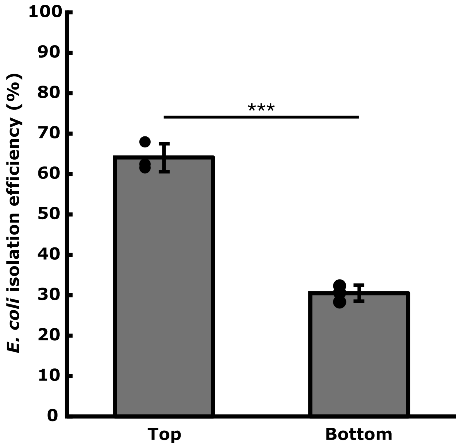

**Supplementary Figure 4: Distribution of bacteria in supernatant and pellet after Smart centrifugation.** *** indicates significance levels p $\leq$ 0.001.

#### Volume reduction with and without diluted percoll as sedimentation interface

Placing a bottom layer of diluted percoll (1:2; percoll:BCM) prevents the aggregation of cells and lysate, resulting in more stable and reproducible flow rates. Supplementary Figure 5 shows how the filtrate and retentate flow in the presence or absence of high-density media (n=5). The use of high-density media in the volume reduction step resulted in better resuspension of the lysate and cells, thus ensuring smoother and more reproducible flow. Additionally, we tested how the presence of a density media at the bottom affected the bacteria isolation efficiency during volume reduction. The isolation efficiency with percoll was found 15% higher than without the percoll, although this difference was not significant 6.

A.

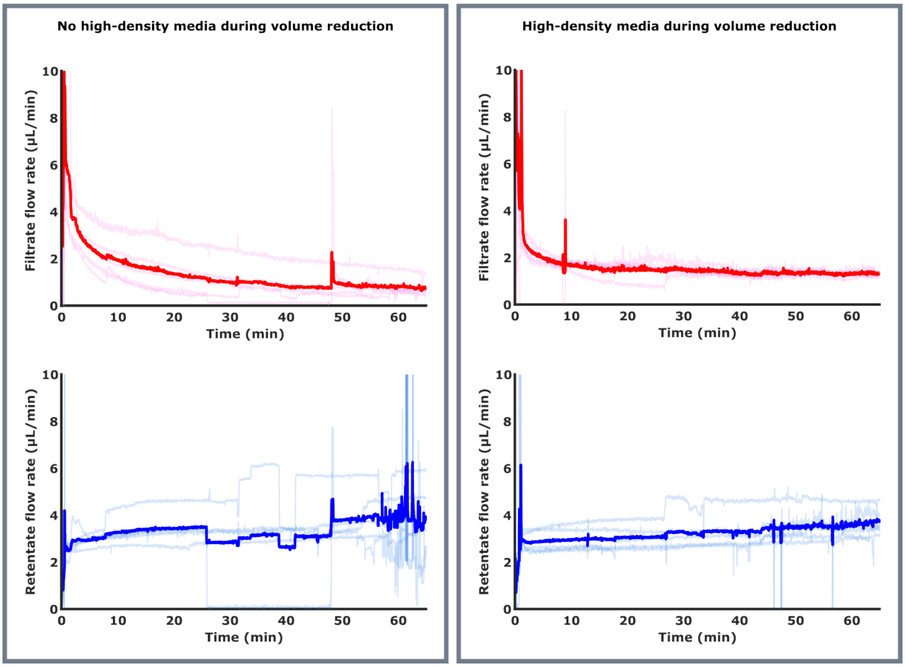

**Supplementary Figure 5**: Filtrate and retentate flows are smoother and more reproducible when using high-density media in the volume reduction step.

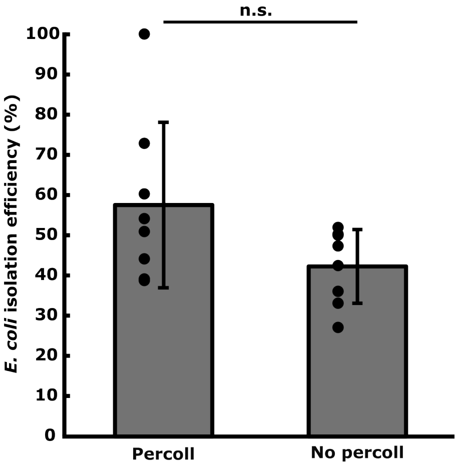

**Supplementary Figure 6**: Comparison of *E. coli* isolation efficiency after volume reduction with and without diluted percoll at the bottom. ns means not significant.

#### Filtrate flow rate and overall trapping efficiency

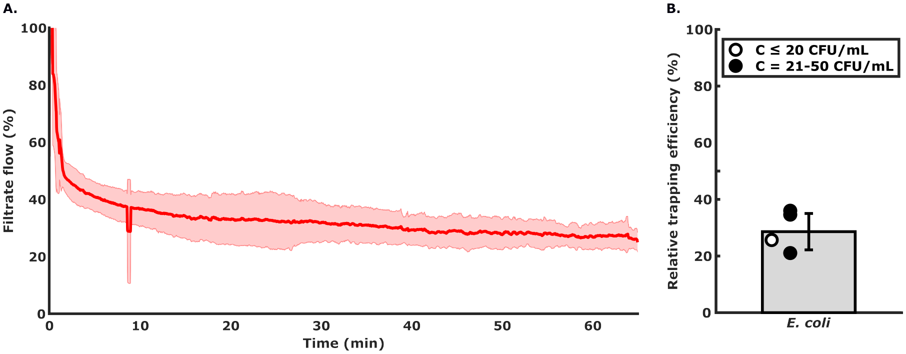

**Supplementary Figure 7**: **A. Filtrate flow rate** for *E. coli*-spiked sample, where the solid line is the mean flow through the microtraps relative to the sample input flow and the shaded area indicates one standard deviation (n=5). **B. Trapping efficiency** for *E. coli*-spiked sample, which is the number of bacteria-containing traps after 70 min relative to the colony-forming units in the sample after smart centrifugation (bar height is mean, error bar is sd).

#### Overall detection rates

The measurements plotted in Supplementary Figure 8 are the same as in Supplementary Figure 4, but now shown as detection rate, meaning the number of bacteria-containing traps after 70 min on-chip sample flow relative to the number of colony-forming units spiked in the blood.

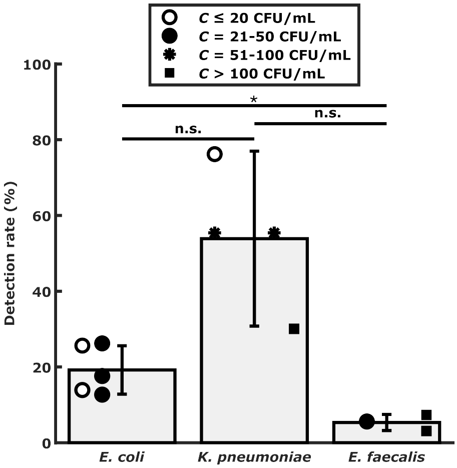

**Supplementary Figure 8**: Detection rate, meaning the number of bacteria-containing traps after 70 min on-chip sample flow relative to the number of colony-forming units spiked in the blood. ns and * indicate significance levels $p > 0.05$ and $p \leq 0.05$, respectivey.

### Comparison table of different bacterial isolation methods

**Supplementary Table 3**: Key performance parameters of smart centrifugation and other separation methods.

| | Bacteria isolation efficiency | RBC removal efficiency | Blood processed (ml) | Time (min) | Throughput (µl/min) | Minimum concentration $C$ detected (CFU/ml) | Volume concentration (Vf / Vi) |
|---|---|---|---|---|---|---|---|
| **Smart centrifugation** | 64% | 99.98% | 2.25 | 5 | 450 | 9 | 1.1 |
| Filter based centrifugation [29] | 26-40% | 99.4% | 1 | 60 | 16.7 | 10 | 18 |
| Compact Disk [30] | 40% | 94% | 7 | 1 | 7000 | 10 | 0.6 |
| Dextran sedimentation [31] | 50-60% | >90% | 10 | 30 | 333 | 10-100 | 0.4 |
| Elastoinertial Separation [28] | >65% | 100% | 1 | 40 | 25 | 40 CFU | 1.5 |
| Inertial lift forces [61] | >80% | 90% | 0.15 | 4 | 37.5 | $>10^5$ | 50 |
| Dielectrophoreis [62] | 87.2% | 100% | 0.05 | 75 | 0.67 | 10000 | 0.01 |
| SAW [63] | 88% | 99.6% | 0.012 | 600 | 0.02 | $4.4.10^7$ | 25 |
| Acoustophoresis [38] | 75% | 99.99% | 1 | 50 | 20 | $5.10^8$ | 3 |
| Magnetic Bead Separation [36] | >80% | >99.99% | 5 | 60 | 83.3 | 1 | 0.2 |

#### Data cleaning and label adjustments

A total of six positions from three experiments were discarded (due to the cropping preprocessing made an error (the barcodes were not clearly visible due to illumination and focus issues). After performing the initial labeling using the developed labeling software, we trained networks and inspected network misclassifications. From this, we corrected the labels of 159 out of 532,782 negative frames to positive and 3 out of 3,231 positive frames to negative in the test set. Furthermore, we corrected the labels of 502 out of 268,105 negative frames to positive and 231 out of 179,811 positive frames to negative in the training set. Only clearly erroneous labels were changed, where we could observe growth, or the traps were completely empty. The labels of ambiguous traps were left unchanged.

#### Video classification

The detection was posed as a video classification problem, where one test instance contained 1-8 frames corresponding to the evaluation time of 0-70 minutes. The number of positive traps changes over time as cells get trapped in previously empty traps, as seen in Table 4. Because of the low concentration of bacteria, only 1% of the traps are positive at 70 minutes.

| **Timestep (minutes)** | **Number of positive traps** | **Fraction positive** |
|---:|---:|---:|
| 0 | 48 | 0.000718 |
| 10 | 152 | 0.002273 |
| 20 | 259 | 0.003873 |
| 30 | 385 | 0.005758 |
| 40 | 428 | 0.006401 |
| 50 | 523 | 0.007821 |
| 60 | 675 | 0.010094 |
| 70 | 721 | 0.010782 |

**Supplementary Table 4**: The test set with the number of positive traps recorded at different time steps for all eight experiments. The whole test set contains 66869 traps, with 66148 being empty at 70 minutes.

The training set consisted of 57,608 time-lapses from eight experiments, containing 5-11 frames, outlined in Table 6, and the test set consisted of 66,869 time-lapses also from eight experiments show in Table 5.

| | Time step (minutes) | | | | | | | |
|---|---|---|---|---|---|---|---|---|
| **Experiment** | 0 | 10 | 20 | 30 | 40 | 50 | 60 | 70 |
| *E. coli* (EXP-25-CA6231) | 9 | 13 | 13 | 13 | 14 | 14 | 14 | 14 |
| *E. faecalis* (EXP-23-CA6223) | 0 | 1 | 1 | 1 | 3 | 6 | 9 | 10 |
| *E. faecalis* (EXP-23-CA6225) | 8 | 33 | 62 | 90 | 105 | 124 | 149 | 168 |
| *E. faecalis* (EXP-23-CA6226) | 2 | 3 | 8 | 10 | 10 | 10 | 10 | 10 |
| *K. pneumoniae* (EXP-23-CA6217) | 21 | 70 | 118 | 189 | 202 | 255 | 354 | 374 |
| *K. pneumoniae* (EXP-23-CA6219) | 2 | 5 | 9 | 18 | 22 | 37 | 58 | 63 |
| *K. pneumoniae* (EXP-23-CA6222) | 0 | 1 | 3 | 5 | 8 | 12 | 14 | 15 |
| *K. pneumoniae* (EXP-23-CA6230) | 6 | 26 | 45 | 59 | 64 | 65 | 67 | 67 |
| **Total** | 48 | 152 | 259 | 385 | 428 | 523 | 675 | 721 |

**Supplementary Table 5**: The test set with the number of positive traps recorded at different time steps for all experiments. The whole test set contains 66869 traps, with 66148 being empty at 70 minutes.

| **Experiment** | **Empty** | **Bacteria** | **Total** |
|---|---|---|---|
| *E. coli* (EXP-24-CA6233) (7 frames) | 7557 | 800 | 8357 |
| *E. coli* (EXP-25-CA6232) (7 frames) | 728 | 3587 | 4315 |
| *E. faecalis* (EXP-25-CA6237) (6 frames) | 3345 | 4870 | 8215 |
| *E. faecalis* (EXP-25-CA6238) (8 frames) | 8362 | 2 | 8364 |
| *K. pneumoniae* (EXP-25-CA6234) (7 frames) | 1942 | 6407 | 8349 |
| *K. pneumoniae* (EXP-25-CA6235) (7 frames) | 671 | 7712 | 8383 |
| No bacteria (EXP-24-CA6229) (11 frames) | 8155 | 0 | 8155 |
| No bacteria (EXP-25-CA6236) (5 frames) | 3470 | 0 | 3470 |
| **Total** | 34230 | 23378 | 57608 |

**Supplementary Table 6**: The training set, showing individual experiments, species, number of frames, positive, negative, and total instances. A trap is shown as empty in the table if none of the frames in the time-lapse contains bacteria.

When employing the image classification models (ResNet 18, DinoV2, and EfficientNet B2), frames were concatenated horizontally to a time-lapse image, allowing us to employ image classification models for the video detection task. When testing (or training) on time-lapses with less than eight frames, the images were padded on the right to 336-pixel width, so all inputs had the same spatial shape (this can also be viewed as masking out later frames). The appearance of image model inputs at different time points is shown in Fig 9A. Furthermore, the performance of all models using potential lower-resolution microscopy was tested by training and testing on subsampled images shown in 9B.

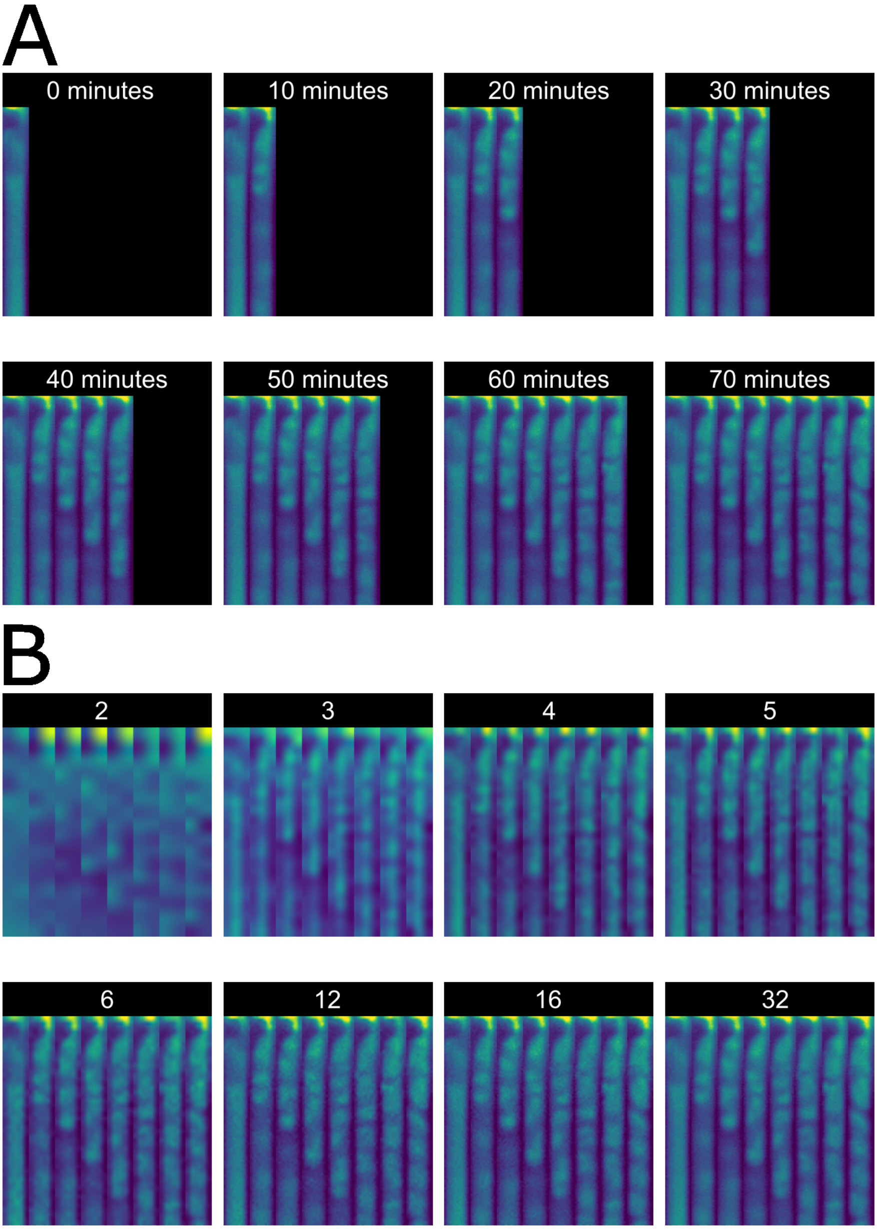

**Supplementary Figure 9**: **Model input** A: Time-lapse images (336x336 pixels) of one trap processed by the image classification models ResNet 18, DinoV2, and EfficientNet B2 during the time evaluation. B: The same trap at 70 minutes imaged at progressively smaller subsampling widths, upsampled to the original size using bicubic interpolation.

Although DinoV2 achieved the best results after 70 minutes, the other models had a slight edge in the first frames, especially the Video ResNet, shown in Fig 10. This may be attributed to the rather unnatural image of one 336x42 frame zero-padded on the right that was inputted to the image networks. However, in later stages, the Video ResNet has poorer generalization. The long timestep of 10 minutes between frames is perhaps not ideal to be processed by temporal convolutions (consecutive frames have too large a difference).

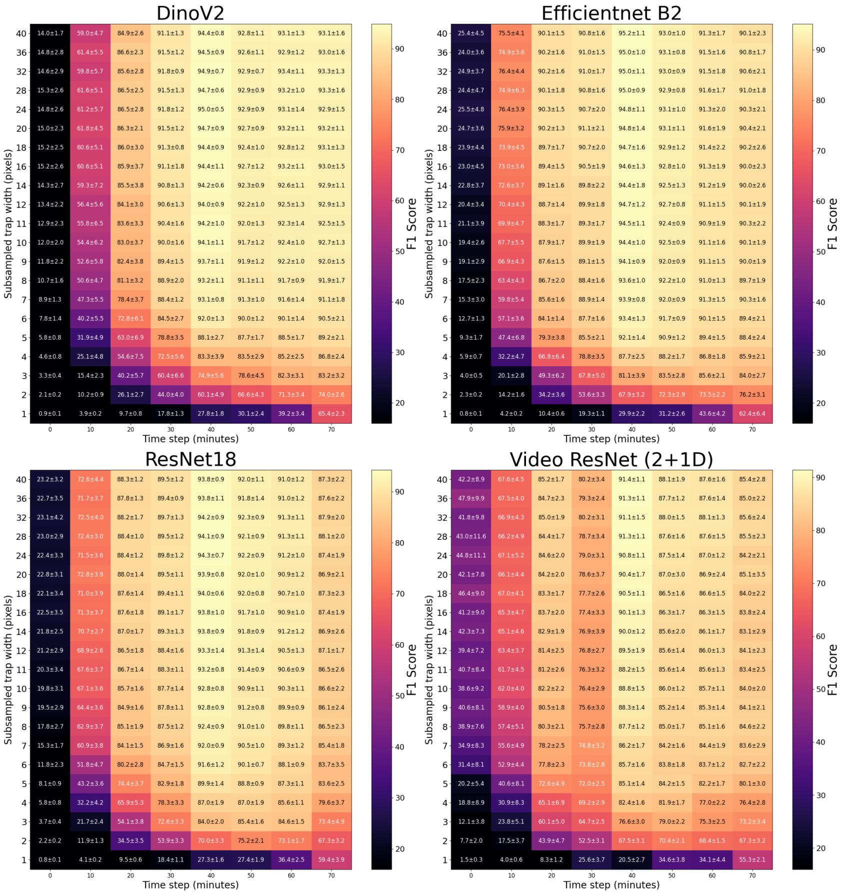

**Supplementary Figure 10**: Comparison of F1 score of all models, framing the problem as video classification, subsampling, and testing on more frames.

#### Misclassifications

The best-achieving model, DinoV2, had 84 false negatives and four false positives from the test partition at 70 minutes. The false positives and a subset of the false negatives are shown in Fig 11. Despite our data cleaning effort, sample a, c, d, e, f have the wrong label and are true negatives (the dark cell-looking objects in a, d and e are actually debris and lack the characteristic "gleaming" bacteria texture)

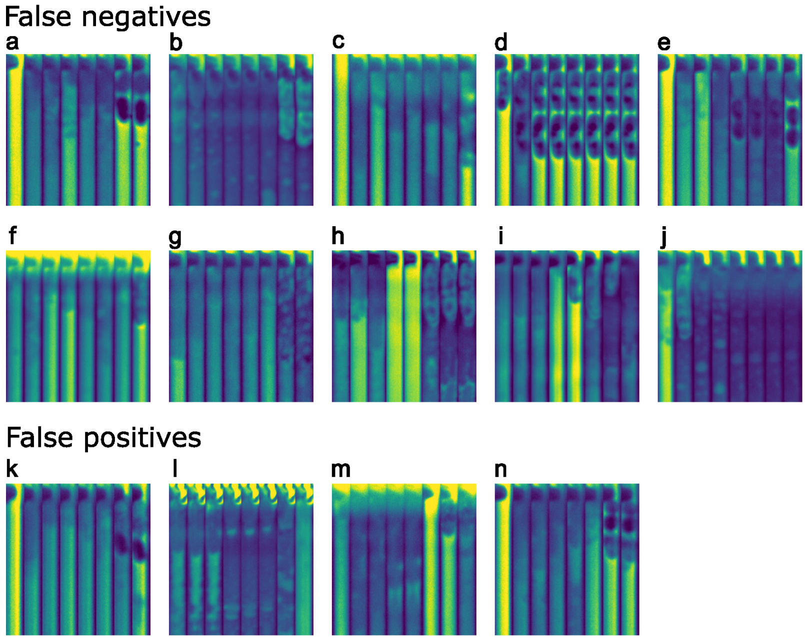

**Supplementary Figure 11**: False negatives and false positives misclassified by DinoV2 at 70 minutes, full resolution.

#### Inference times and computational complexity

The FLOPs count stated in the results section was calculated on inference shapes of 336x336 pixels for the image models and 8x336x42 pixels for Video ResNet. The inference latency of one sample on the A100 GPU was $1.410 \pm 0.086$ms for ResNet 18, $1.693 \pm 0.051$ms for EfficientNet B2, $3.085 \pm 0.032$ms for DinoV2, and $1.573 \pm 0.046$ms for Video ResNet (2+1D) (using Pytorch data loader with 5 workers, test batch size 32). However, the inference times could be improved using software and hardware optimizations in a live setting.

#### Posing the problem as single-frame image classification

In our study, we posed the problem as a video classification problem, treating a time-lapse of an individual trap as one classification instance. A trap was assigned a positive label if it contained or had previously contained any cells until the evaluation time point. This problem framing aligns better with a potential clinical application. However, since we have frame-level annotations, we can treat it as a simple image classification problem and independently classify each frame, disregarding the time lapses. One classification instance is then a single frame from any trap time-lapse in any experiment and has a positive label only if it contains bacteria. Processing single-frames could be more feasible if operating on resource-constrained hardware. The smaller memory footprint allowed us to use the whole 1300x40 pixel trap images as inputs as opposed to cropping the height at 336 pixels for the video classification. The train and test sets are shown in table 7 and table 8. Analogous to the video classification, all single-frame networks were retrained 30 times with unique random seeds for reproducibility and statistics. When performing subsampling experiments, the images were upsampled back to the original size using bicubic interpolation before feeding into the networks. All single-frame networks were trained for 50 epochs using batch size 256, a learning rate of $10^{-3}$ for all CNN:s and $10^{-5}$ for DinoV2, otherwise the same settings as when training for video classification (except random frame erase).

| **Experiment** | **Empty** | **Bacteria** | **Total** |
|---|---|---|---|
| E. coli (EXP-24-CA6233) (7 frames) | 52467 | 6154 | 58621 |
| E. coli (EXP-25-CA6232) (7 frames) | 5089 | 42832 | 47921 |
| E. faecalis (EXP-25-CA6237) (6 frames) | 20013 | 29503 | 49516 |
| E. faecalis (EXP-25-CA6238) (8 frames) | 66928 | 100 | 67028 |
| K. pneumoniae (EXP-25-CA6234) (7 frames) | 11325 | 47266 | 58591 |
| K. pneumoniae (EXP-25-CA6235) (7 frames) | 4478 | 54227 | 58705 |
| No bacteria (EXP-24-CA6229) (11 frames) | 90184 | 0 | 90184 |
| No bacteria (EXP-25-CA6236) (5 frames) | 17350 | 0 | 17350 |
| Total | 267834 | 180082 | 447916 |

**Supplementary Table 7**: The training set for single-frame classification, showing individual experiments, species, positive, negative, and total single-frame images.

#### Subsampling single-frames

We also wanted to study the performance at lower resolution for the single-frames, training, and testing on subsampled images. The results shown in Fig 12, a sharp perfomance drop happens in about 8-pixel width, indicating that there is plenty of room to use lower-resolution microscopy for the task.

| **Experiment** | **Empty** | **Bacteria** | **Total** |
|---|---|---|---|
| E. coli (EXP-25-CA6231) | 67034 | 144 | 67178 |
| E. faecalis (EXP-23-CA6223) | 66505 | 23 | 66528 |
| E. faecalis (EXP-23-CA6225) | 66413 | 751 | 67164 |
| E. faecalis (EXP-23-CA6226) | 67133 | 63 | 67196 |
| K. pneumoniae (EXP-23-CA6217) | 65164 | 1728 | 66892 |
| K. pneumoniae (EXP-23-CA6219) | 66642 | 214 | 66856 |
| K. pneumoniae (EXP-23-CA6222) | 67140 | 58 | 67198 |
| K. pneumoniae (EXP-23-CA6230) | 66595 | 406 | 67001 |
| Total | 532626 | 3387 | 536013 |

**Supplementary Table 8**: The test set for single-frame classification, showing individual experiments, species, positive, negative, and total single-frame images.

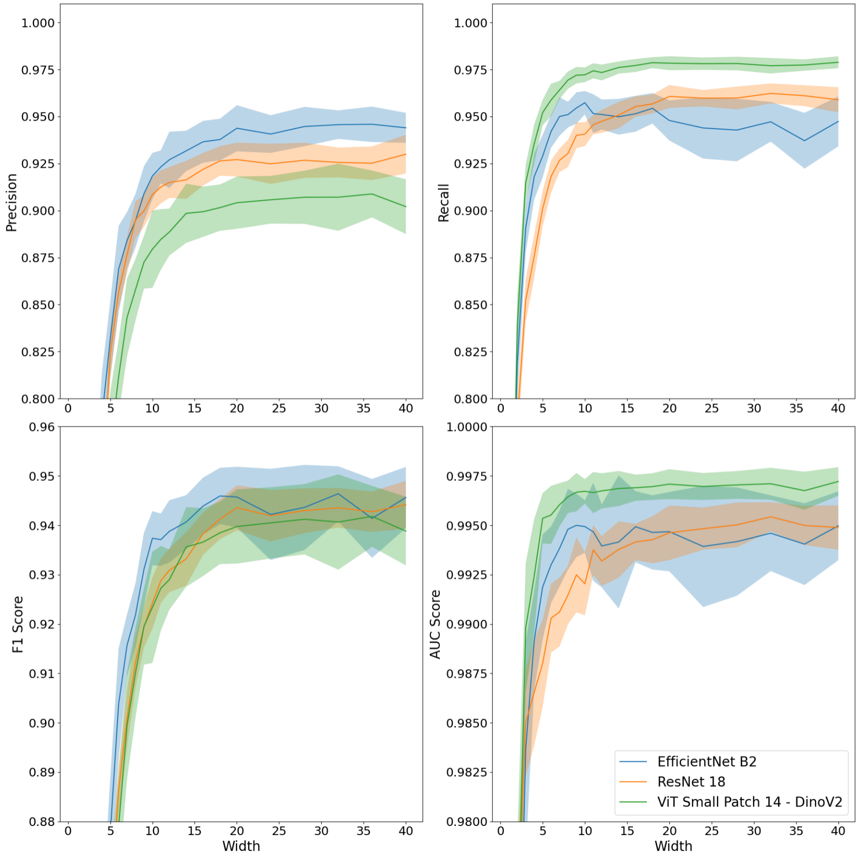

**Supplementary Figure 12**: Performance metrics of all image models when processing subsampled images. Lines show mean, and shaded areas show standard deviations among the 30 retrainings.

#### Images of cells at different time points

Typically, frames from earlier timesteps were more challenging to classify because the cells are significantly smaller, lack the characteristic bacteria cell texture, and are situated at the very top of the microfluidic trap. Therefore, to study the performance of the single-frame classification network at different time points, we partitioned the test set based on the time at which each frame was captured and calculated the classification performance for each subset. The interesting results are shown in Fig 13. For the first few frames, it is more advantageous to use single-frame models. When processing 1300x40 frames, it is easier for the network to see the top end of the bacterial column, which could be important at lower resolution. The results at 70 minutes were 95.3+-0.6 for DinoV2, 95.2+-0.6 for Efficientnet B2, and 94.7+-0.5 for ResNet 18. Note that it is problematic to compare the final performance to the video models: Hard samples where traps previously contained cells that subsequently escaped are still labeled positive for the video models at 70 minutes.

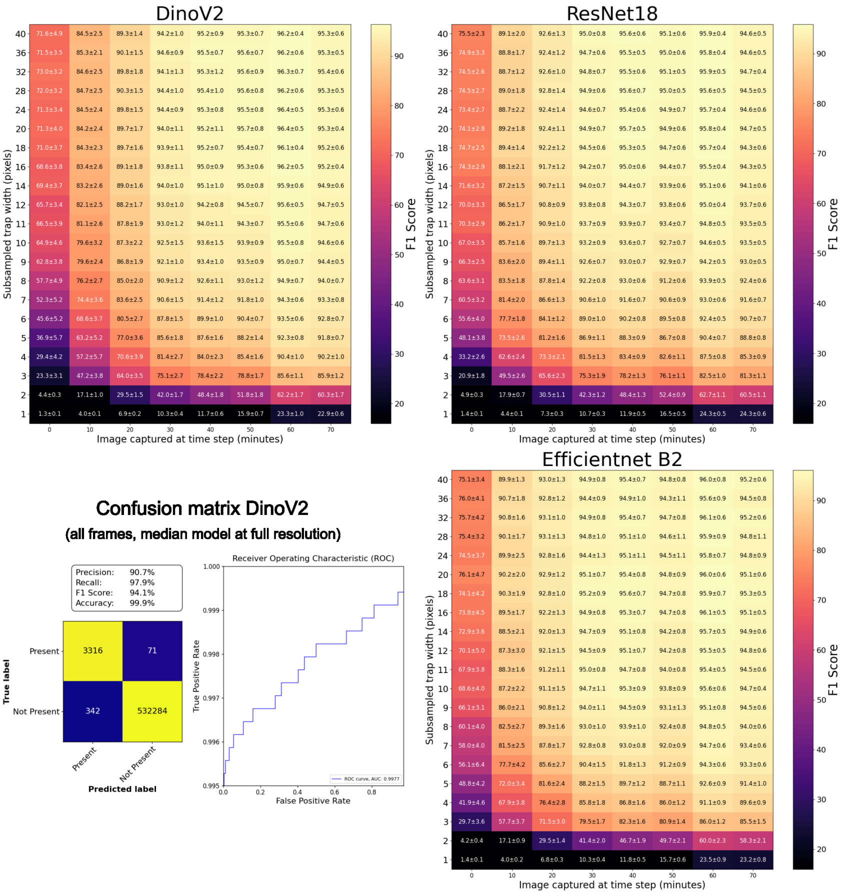

**Supplementary Figure 13**: Single-frame classification results, partitioning the test set according to when a test frame was taken. The lower left shows the confusion matrix and AUC of the best model, DinoV2, testing on frames from all timesteps, showing the model with the median F1-score of the 30 retrainings.

#### Reshaping the oblong images during single-frame classification

All our studies used networks initialized with pretrained weights, either on ImageNet, or self-supervised DinoV2. The pretraining was performed on natural images with image size 224x224 pixels, very different from our oblong 1300x40 pixel trap images. We wanted to see how reshaping affects performance, so we performed experiments split up the trap into patches of 224x40 and stacked horizontally using three strategies:

1. Reshaping and performing our usual padding to 42-pixel width, resulting in an input image of 224x256.
2. Reshaping but cropping each trap to 39 pixels and cropping the final reshaped frame 10 pixels to an image size of 224x224.
3. Performing no reshaping, using the unchanged 1300x40 pixel images.

Results are shown in Fig 14, each network was retrained 50 times using different random seeds. Despite ResNets and EfficientNets using the same Global Average Pooling in the final layer, EfficientNet performs better. Thus, it is better at extracting features regardless of shape. For all our experiments, we chose the first strategy, thus reshaping the single-frame images 224x256 before feeding into the network.

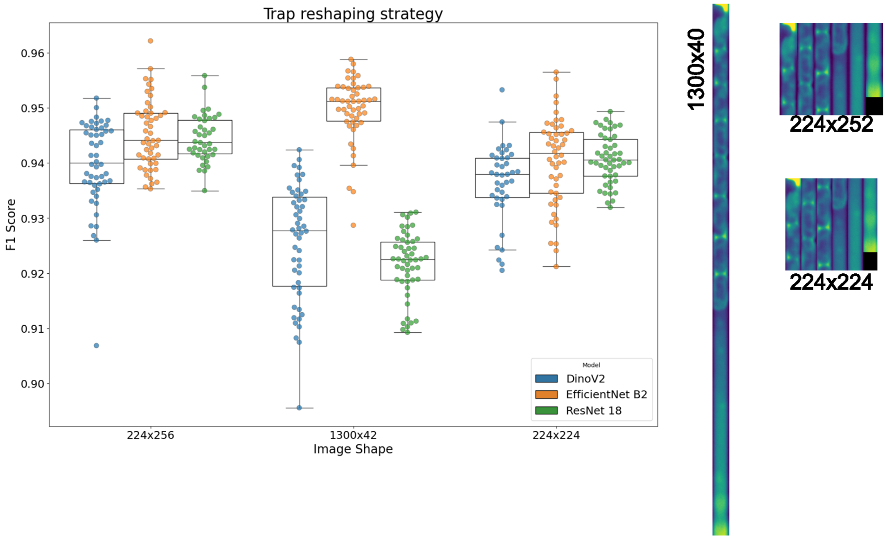

**Supplementary Figure 14**: Retraining networks using different reshaping strategies.

#### Image preprocessing and cropping out traps

An image processing pipeline was further improved from [50] to rotate, crop out, and stabilize the individual traps in the microfluidic chip. The preprocessing code is available in the released software package [59] and was developed using OpenCV[64], Numpy[65], Scipy[66], and MatPlotLib[67]. The pipeline consists of the following steps:

1. Rotate the frames by a few degrees to align the chip structure horizontally.
2. Project the image on the horizontal axis to find the location of the traps and barcodes in each frame.
3. Determine which peaks correspond to barcodes and assign a location ID to each trap.

4. Identify the location of the physical stop protruding out from the trap wall and determine where to crop the traps along the vertical axis.
5. Crop and stack the 1300x40 pixel trap images, and perform fine-grained stabilization using StackReg.[68]

For the rotation step, we first applied Gaussian blur to the images, then adaptive thresholding, calculating the contours, and finally projecting the binary image on the horizontal and vertical axes. When the chip structure was perfectly horizontal, the sums of the projections were at maximum values (trap walls aligned vertically). A grid search was performed with progressively smaller steps to find the optimal rotation angle.

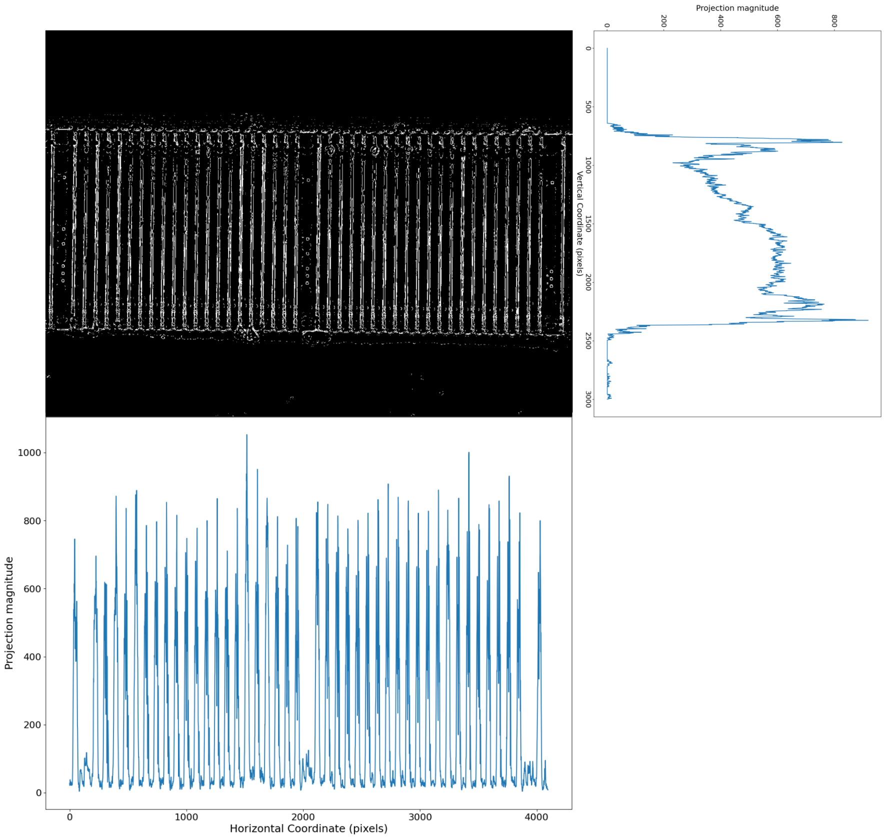

**Supplementary Figure 15**: Projections on the horizontal and vertical axes were used to find the optimal rotation angle.

Then, the raw images were projected on the horizontal axis, followed by 1D Gaussian blur, revealing the horizontal location of the traps 16A. The barcodes were also visible as peaks in the projection. The traps were cropped along the vertical axis by a 20-pixel displacement on each side of a peak. A barcode score was assigned to the three leftmost, rightmost, and middle crop-outs to determine the most likely barcode locations. This score was based on whether there were any peaks when projecting the cropped-out images on the vertical axis, the 75th percentile pixel value (barcodes images had darker pixels on average), and a few other metrics. In practice, finding a robust metric was challenging: a trap could be very dark with bright peaks if filled with cocci cells, and conversely, a barcode could be bright if it contained many barcode dots. By knowing the barcode locations, we could assign a trap ID to each crop out and pair them across the different frames in the time-lapse. This was necessary as the raw images often exhibited significant drift during the time-lapse.

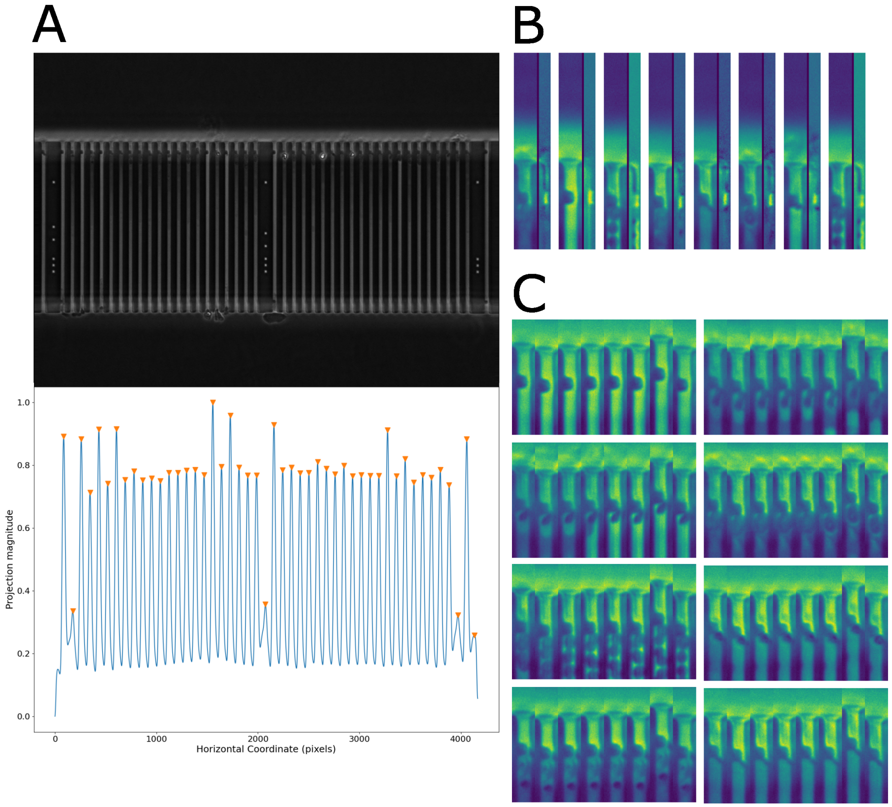

**Supplementary Figure 16**: A: Projecting the traps on the horizontal axis followed by applying 1D Gaussian 1D kernel, with the peaks showing the trap locations. B: Using the asymmetry introduced by the physical stop protruding out from the side of the wall to find the location to crop horizontally. Images of eight separate traps are shown, horizontally concatenated with a dark border and the calculated image difference, revealing the stop as a bright spot. C: The fine-grained stabilization of the cropped-out phase-contrast time-lapse stack was calculated using the movement of the physical stop-stop (300x40 pixel crop). The images show eight different traps and the corresponding frames in the time-lapse horizontally concatenated. There was a larger vertical shift in the seventh frame, later corrected with the fine-grained stabilization.

The vertically cropped trap images were then cropped on the horizontal axis right at the location of the physical stop. The stop introduced an asymmetry in the image, and when calculating the image difference of the left and the horizontally flipped right image-half (cropped vertically at the center) revealed the stop as a bright peak 16B, pseudocode outlined below:

$$
\begin{aligned}
&\text{Let } \mathbf{IM}_{\text{trap}} \in \mathbb{R}^{1300\times 40} \\
&\mathbf{IM}_{\text{left}} = \mathbf{IM}_{\text{trap}}[:, 1:20], \\
&\mathbf{IM}_{\text{right}} = \mathbf{IM}_{\text{trap}}[:, 21:40], \\
&\mathbf{IM}_{\text{diff}} = \text{FlipLeftRight}(\mathbf{IM}_{\text{right}}) - \mathbf{IM}_{\text{left}}.
\end{aligned}
$$

After cropping, the time-lapse of each trap was fine-grained stabilized using StackReg[68] using only translation. The transformation matrices were calculated using a small 300x40 pixel cropout containing the physical stop, a few samples are shown in 16C. Finally, the color map Vidiris[53] was applied to the single-channel phase-contrast microscopy images to be easily viewed by a human labeler.

#### Tabled experimental measurements and their results

The tables below contain the detailed experimental description and raw data of plate counts for all the experiments included in this study.

Expt. id is the unique identification number of the experiment.

Expt. type is the type of experiment, meaning either coventional centrifugation, smart centrifugation, volume reduction and detection in microfluidics trap, refer Methods.

Control plate count is the bacteria count after plating 100 µl of the spiking solution used to spike the blood (n=3). Columns 2-4 show the number of colonies on the plate. The last column shows the mean count.

Expt. plate count is the bacteria count after plating 100 µl of recovered solution after a centrifugation steps. Columns 2-4 show the number of colonies on the plate. The last column shows the mean count.

Volume recovered (ml) is the volume recovered after centrifugation.

Isolation efficiency (%) is the efficiency of bacterial separation of the experiment, meaning the number of bacteria recovered relative to the number of bacteria initially spiked.

Concentration in blood (CFU/ml) is the concentration of bacteria in the spiked blood, $C$.

Bacteria (t60), (t70) and (tend) are the number of bacteria trapped in the microtraps after 60 min on-chip processing, 70 min on-chip processing, and at the end of the experiment.

Overall assay performance is the number of positive microtraps detected, N, versus the bacterial concentration in the blood sample, C.

**Supplementary Table 9**: Raw data of plate counts of all experiments with *E. coli*.

| | | | | Mean |
|---|---|---|---|---|
| Expt. id (Date / Name) | 2022-12-02 | | | |
| Expt. type | Conventional centrifugation | | | |
| Control plate count | 42 | 71 | 69 | 60.7 |
| Expt. plate count | 23 | 50 | 55 | 42.7 |
| Volume recovered (ml) | 2.0 | | | |
| Isolation efficiency (%) | 39.7 | | | |
| Concentration in blood (CFU/ml) | 537.7 | | | |
| Expt. id (Date / Name) | 2022-12-05 | | | |
| Expt. type | Conventional centrifugation | | | |
| Control plate count | 48 | 32 | 29 | 36 |
| Expt. plate count | 27 | 14 | 26 | 22.3 |
| Volume recovered (ml) | 1.9 | | | |

| | | | | |
|---|---|---|---|---|
| Isolation efficiency (%) | 32.9 | | | |
| Concentration in blood (CFU/ml) | 322.0 | | | |
| Expt. id (Date / Name) | 2022-12-07 | | | |
| Expt. type | Conventional centrifugation | | | |
| Control plate count | 45 | 32 | | 38.5 |
| Expt. plate count | 12 | 21 | 26 | 19.7 |
| Volume recovered (ml) | 1.9 | | | |
| Isolation efficiency (%) | 26.7 | | | |
| Concentration in blood (CFU/ml) | 295.8 | | | |
| Expt. id (Date / Name) | 2023-03-02 (a) | | | |
| Expt. type | Smart centrifugation | | | |
| Control plate count | 58 | 52 | 43 | 51 |
| Expt. plate count | 42 | 36 | 39 | 39 |
| Volume recovered (ml) | 2.8 | | | |
| Isolation efficiency (%) | 71.4 | | | |
| Concentration in blood (CFU/ml) | 680 | | | |
| Expt. id (Date / Name) | 2023-03-02 (b) | | | |
| Expt. type | Smart centrifugation | | | |
| Control plate count | 58 | 52 | 43 | 51 |
| Expt. plate count | 41 | 30 | | 35.5 |
| Volume recovered (ml) | 2.7 | | | |
| Isolation efficiency (%) | 62.6 | | | |
| Concentration in blood (CFU/ml) | 680 | | | |
| Expt. id (Date / Name) | 2023-03-02 (c) | | | |
| Expt. type | Smart centrifugation | | | |
| Control plate count | 75 | 53 | 70 | 66 |
| Expt. plate count | 48 | 42 | | 45 |
| Volume recovered (ml) | 3.0 | | | |
| Isolation efficiency (%) | 67.4 | | | |
| Concentration in blood (CFU/ml) | 880 | | | |
| Expt. id (Date / Name) | 2023-07-05 (a) | | | |

| | | | | |
|---|---|---|---|---|
| Expt. type | Smart centrifugation | | | |
| Control plate count | 161 | 143 | 158 | 154.0 |
| Expt. plate count | 81 | 75 | 78 | 78.0 |
| Volume recovered (ml) | 3.1 | | | |
| Isolation efficiency (%) | 79.8 | | | |
| Concentration in blood (CFU/ml) | 1346 | | | |
| Expt. id (Date / Name) | 2023-07-05 (b) | | | |
| Expt. type | Smart centrifugation | | | |
| Control plate count | 161 | 143 | 158 | 154.0 |
| Expt. plate count | 75 | 64 | 97 | 78.7 |
| Volume recovered (ml) | 3.1 | | | |
| Isolation efficiency (%) | 80.5 | | | |
| Concentration in blood (CFU/ml) | 1346 | | | |
| Expt. id (Date / Name) | EC Chip Trial 3 | | | |
| Expt. type | Smart centrifugation + Microfluidic detection | | | |
| Control plate count | 101 | 108 | 99 | 102.7 |
| Expt. plate count | 10 | 9 | 4 | 7.7 |
| Volume recovered (ml) | 2.5 | | | |
| Isolation efficiency (%) | 29.9 | | | |
| Concentration in blood (CFU/ml) | 51.0 | | | |
| Bacteria (t60) | 3 | | | |
| Bacteria (tend) | 5 | | | |
| Overall assay performance (%) | 17.4 | | | |
| Expt. id (Date / Name) | EC Chip Trial 1 | | | |
| Expt. type | Smart centrifugation + Microfluidic detection | | | |
| Control plate count | 82 | 71 | 74 | 75.7 |
| Expt. plate count | 11 | 9 | 6 | 8.7 |
| Volume recovered (ml) | 2.5 | | | |
| Isolation efficiency (%) | 50.9 | | | |
| Concentration in blood (CFU/ml) | 38 | | | |
| Bacteria (t60) | 14 | | | |

| | | | | |
|---|---|---|---|---|
| Bacteria (tend) | 19 | | | |
| Overall assay performance (%) | 43.8 | | | |
| Expt. id (Date / Name) | EC Chip Trial 2 | | | |
| Expt. type | Smart centrifugation + Microfluidic detection | | | |
| Control plate count | 36 | 42 | 48 | 42.0 |
| Expt. plate count | 7 | 7 | 5 | 6.3 |
| Volume recovered (ml) | 2.5 | | | |
| Isolation efficiency (%) | 60.3 | | | |
| Concentration in blood (CFU/ml) | 21 | | | |
| Bacteria (t60) | 4 | | | |
| Bacteria (tend) | 6 | | | |
| Overall assay performance (%) | 21.0 | | | |
| Expt. id (Date / Name) | EC Chip Trial 6 | | | |
| Expt. type | Smart centrifugation + Microfluidic detection | | | |
| Control plate count | 35 | 50 | 37 | 40.7 |
| Expt. plate count | 9 | 8 | 3 | 6.7 |
| Volume recovered (ml) | 2.5 | | | |
| Isolation efficiency (%) | 72.9 | | | |
| Concentration in blood (CFU/ml) | 20 | | | |
| Bacteria (t60) | 9 | | | |
| Bacteria (tend) | 24 | | | |
| Overall assay performance (%) | 52.5 | | | |
| Expt. id (Date / Name) | EC Chip Trial 5 | | | |
| Expt. type | Smart centrifugation + Microfluidic detection | | | |
| Control plate count | 38 | 35 | 42 | 38.3 |
| Expt. plate count | 6 | 6 | 2 | 4.7 |
| Volume recovered (ml) | 2.5 | | | |
| Isolation efficiency (%) | 54.1 | | | |

| | | | | |
|---|---|---|---|---|
| Concentration in blood (CFU/ml) | 19 | | | |
| Bacteria (t60) | 6 | | | |
| Bacteria (tend) | 8 | | | |
| Overall assay performance (%) | 18.6 | | | |
| Expt. id (Date / Name) | EC Chip Trial 4 | | | |
| Expt. type | Smart centrifugation + Microfluidic detection | | | |
| Control plate count | 17 | 23 | 12 | 17.3 |
| Expt. plate count | 6 | 4 | 3 | 4.3 |
| Volume recovered (ml) | 2.5 | | | |
| Isolation efficiency (%) | 100.0 | | | |
| Concentration in blood (CFU/ml) | 9.0 | | | |
| Bacteria (t60) | 5.0 | | | |
| Bacteria (tend) | 8.0 | | | |
| Overall assay performance (%) | 41.0 | | | |
| Expt. id (Date / Name) | 2024-02-22 (a) | | | |
| Expt.desciption | Smart centrifugation + Standard Up-concentration | | | |
| Control plate count | 128 | 126 | 145 | 133.0 |
| Expt. plate count | 114 | 137 | 119 | 123.3 |
| Volume recovered (ml) | 0.6 | | | |
| Isolation efficiency (%) | 44.2 | | | |
| Concentration in blood (CFU/ml) | 682 | | | |
| Expt. id (Date / Name) | 2024-02-22 (b) | | | |
| Expt.desciption | Smart centrifugation + Standard Up-concentration | | | |
| Control plate count | 128 | 126 | 145 | 133.0 |
| Expt. plate count | 120 | 100 | 104 | 108.0 |
| Volume recovered (ml) | 0.6 | | | |
| Isolation efficiency (%) | 38.7 | | | |
| Concentration in blood (CFU/ml) | 682 | | | |

| | | | | |
|---|---|---|---|---|
| Expt. id (Date / Name) | 2024-02-22 (c) | | | |
| Expt.desciption | Smart centrifugation + Standard Up-concentration | | | |
| Control plate count | 128 | 126 | 145 | 133.0 |
| Expt. plate count | 120 | 105 | 103 | 109.3 |
| Volume recovered (ml) | 0.6 | | | |
| Isolation efficiency (%) | 44.2 | | | |
| Concentration in blood (CFU/ml) | 682 | | | |
| Expt. id (Date / Name) | 2024-02-22 (d) | | | |
| Expt.desciption | Smart centrifugation + Standard Up-concentration without percoll | | | |
| Control plate count | 128 | 126 | 145 | 133.0 |
| Expt. plate count | 71 | 85 | 71 | 75.7 |
| Volume recovered (ml) | 0.6 | | | |
| Isolation efficiency (%) | 27.1 | | | |
| Concentration in blood (CFU/ml) | 682 | | | |
| Expt. id (Date / Name) | 2024-02-22 (e) | | | |
| Expt.desciption | Smart centrifugation + Standard Up-concentration without percoll | | | |
| Control plate count | 128 | 126 | 145 | 133 |
| Expt. plate count | 99 | 81 | 97 | 92.3 |
| Volume recovered (ml) | 0.6 | | | |
| Isolation efficiency (%) | 33.1 | | | |
| Concentration in blood (CFU/ml) | 682.0 | | | |
| Expt. id (Date / Name) | 2024-02-22 (f) | | | |
| Expt.desciption | Smart centrifugation + Standard Up-concentration without percoll | | | |
| Control plate count | 128 | 126 | 145 | 133.0 |
| Expt. plate count | 101 | 107 | 94 | 100.7 |
| Volume recovered (ml) | 0.6 | | | |

| | | | | |
|---|---|---|---|---|
| Isolation efficiency (%) | 36.1 | | | |
| Concentration in blood (CFU/ml) | 682 | | | |
| Expt. id (Date / Name) | 2024-02-26 (a) | | | |
| Expt.desciption | Smart centrifugation + Standard Up-concentration without percoll | | | |
| Control plate count | 134 | 136 | 133 | 134.3 |
| Expt. plate count | 209 | 229 | 202 | 213.3 |
| Volume recovered (ml) | 0.5 | | | |
| Isolation efficiency (%) | 50.0 | | | |
| Concentration in blood (CFU/ml) | 852.9 | | | |
| Expt. id (Date / Name) | 2024-02-26 (b) | | | |
| Expt.desciption | Smart centrifugation + Standard Up-concentration without percoll | | | |
| Control plate count | 134 | 136 | 133 | 134.3 |
| Expt. plate count | 194 | 199 | 212 | 201.7 |
| Volume recovered (ml) | 0.5 | | | |
| Isolation efficiency (%) | 47.3 | | | |
| Concentration in blood (CFU/ml) | 852.9 | | | |
| Expt. id (Date / Name) | 2024-02-26 (c) | | | |
| Expt.desciption | Smart centrifugation + Standard Up-concentration without percoll | | | |
| Control plate count | 134 | 136 | 133 | 134.3 |
| Expt. plate count | 159 | 195 | 190 | 181.3 |
| Volume recovered (ml) | 0.5 | | | |
| Isolation efficiency (%) | 42.5 | | | |
| Concentration in blood (CFU/ml) | 852.9 | | | |
| Expt. id (Date / Name) | 2024-02-26 (d) | | | |

| | | | | |
|---|---|---|---|---|
| Expt.desciption | Smart centrifugation + Standard Up-concentration without percoll | | | |
| Control plate count | 134 | 136 | 133 | 134.3 |
| Expt. plate count | 198 | 231 | 235 | 221.3 |
| Volume recovered (ml) | 0.5 | | | |
| Isolation efficiency (%) | 51.9 | | | |
| Concentration in blood (CFU/ml) | 852.9 | | | |
| Expt. id (Date / Name) | 2024-02-26 (e) | | | |
| Expt.desciption | Smart centrifugation + Standard Up-concentration without percoll | | | |
| Control plate count | 134 | 136 | 133 | 134.3 |
| Expt. plate count | 222 | 207 | 215 | 214.7 |
| Volume recovered (ml) | 0.5 | | | |
| Isolation efficiency (%) | 50.3 | | | |
| Concentration in blood (CFU/ml) | 852.9 | | | |
| Expt. id (Date / Name) | 2024-02-20 (a) | | | |
| Expt.desciption | Smart centrifugation | | | |
| Control plate count | 112 | 122 | 102 | 112.0 |
| Expt. plate count | 34 | 39 | | 36.5 |
| Volume recovered (ml) | 2.6 | | | |
| Isolation efficiency (%) | 62.5 | | | |
| Concentration in blood (CFU/ml) | 933.3 | | | |
| Expt. id (Date / Name) | 2024-02-20 (b) | | | |
| Expt.desciption | Smart centrifugation | | | |
| Control plate count | 112 | 122 | 102 | 112.0 |
| Expt. plate count | 41 | 31 | | 36 |
| Volume recovered (ml) | 2.6 | | | |
| Isolation efficiency (%) | 61.7 | | | |
| Concentration in blood (CFU/ml) | 933.3 | | | |
| Expt. id (Date / Name) | 2024-02-22 (g) | | | |
| Expt.desciption | Smart centrifugation | | | |

| | | | | |
|---|---|---|---|---|
| Control plate count | 128 | 126 | 145 | 133.0 |
| Expt. plate count | 37 | 40 | 35 | 37.3 |
| Volume recovered (ml) | 2.8 | | | |
| Isolation efficiency (%) | 68.1 | | | |
| Concentration in blood (CFU/ml) | 682 | | | |

**Supplementary Table 10**: Raw data of plate counts of all experiments with *K. pneumoniae.*

| | | | | Mean |
|---|---|---|---|---|
| Expt. id (Date / Name) | 2023-09-22 (a) | | | |
| Expt. type | Smart centrifugation | | | |
| Control plate count | 131 | | | 131.0 |
| Expt. plate count | 123 | 116 | 139 | 126.0 |
| Volume recovered (ml) | 2.7 | | | |
| Isolation efficiency (%) | 95.2 | | | |
| Concentration in blood (CFU/ml) | 1587.8 | | | |
| Expt. id (Date / Name) | 2023-09-22 (b) | | | |
| Expt. type | Smart centrifugation | | | |
| Control plate count | 131 | | | 131.0 |
| Expt. plate count | 118 | 126 | 110 | 118.0 |
| Volume recovered (ml) | 2.7 | | | |
| Isolation efficiency (%) | 89.2 | | | |
| Concentration in blood (CFU/ml) | 1587.8 | | | |
| Expt. id (Date / Name) | 2023-09-22 (c) | | | |
| Expt. type | Smart centrifugation | | | |
| Control plate count | 131 | | | 131.0 |
| Expt. plate count | 21 | 12 | 11 | 14.7 |
| Volume recovered (ml) | 2.6 | | | |
| Isolation efficiency (%) | 98.0 | | | |
| Concentration in blood (CFU/ml) | 172.9 | | | |
| Expt. id (Date / Name) | 2023-09-22 (d) | | | |
| Expt. type | Smart centrifugation | | | |
| Control plate count | 131 | | | 131.0 |
| Expt. plate count | 1 | 0 | 3 | 1.3 |
| Volume recovered (ml) | 2.6 | | | |
| Isolation efficiency (%) | 97.0 | | | |
| Concentration in blood (CFU/ml) | 15.8 | | | |
| Expt. id (Date / Name) | 2023-09-22 (e) | | | |
| Expt. type | Smart centrifugation | | | |
| Control plate count | 131 | | | 131.0 |
| Expt. plate count | 1 | 2 | 2 | 1.7 |

| | | | | |
|---|---|---|---|---|
| Volume recovered (ml) | 2.7 | | | |
| Isolation efficiency (%) | 125.9 | | | |
| Concentration in blood (CFU/ml) | 15.8 | | | |
| Expt. id (Date / Name) | 2023-09-22 (f) | | | |
| Expt. type | Smart centrifugation | | | |
| Control plate count | 131 | | | 131.0 |
| Expt. plate count | 1 | 0 | 3 | 1.3 |
| Volume recovered (ml) | 2.6 | | | |
| Isolation efficiency (%) | 97.0 | | | |
| Concentration in blood (CFU/ml) | 15.9 | | | |
| Expt. id (Date / Name) | 2023-09-22 (g) | | | |
| Expt. type | Smart centrifugation | | | |
| Control plate count | 131 | | | 131.0 |
| Expt. plate count | 1 | 2 | 2 | 1.7 |
| Volume recovered (ml) | 2.7 | | | |
| Isolation efficiency (%) | 125.9 | | | |
| Concentration in blood (CFU/ml) | 15.9 | | | |
| Expt. id (Date / Name) | 2023-09-26 (a) | | | |
| Expt. type | Smart centrifugation | | | |
| Control plate count | 212 | 231 | 213 | 218.7 |
| Expt. plate count | 187 | 184 | 189 | 186.7 |
| Volume recovered (ml) | 2.7 | | | |
| Isolation efficiency (%) | 82.9 | | | |
| Concentration in blood (CFU/ml) | 2650.5 | | | |
| Expt. id (Date / Name) | 2023-09-26 (b) | | | |
| Expt. type | Smart centrifugation | | | |
| Control plate count | 212 | 231 | 213 | 218.7 |
| Expt. plate count | 25 | 18 | 15 | 19.3 |
| Volume recovered (ml) | 2.6 | | | |
| Isolation efficiency (%) | 84.3 | | | |
| Concentration in blood (CFU/ml) | 265 | | | |
| Expt. id (Date / Name) | 2023-09-26 (c) | | | |
| Expt. type | Smart centrifugation | | | |

| | | | | |
|---|---|---|---|---|
| Control plate count | 212 | 231 | 213 | 218.7 |
| Expt. plate count | 17 | 18 | 18 | 17.7 |
| Volume recovered (ml) | 2.6 | | | |
| Isolation efficiency (%) | 77.0 | | | |
| Concentration in blood (CFU/ml) | 265 | | | |
| Expt. id (Date / Name) | 2023-09-27 (a) | | | |
| Expt. type | Smart centrifugation | | | |
| Control plate count | 175 | 186 | 187 | 182.7 |
| Expt. plate count | 26 | 25 | 19 | 23.3 |
| Volume recovered (ml) | 2.2 | | | |
| Isolation efficiency (%) | 103.0 | | | |
| Concentration in blood (CFU/ml) | 221.4 | | | |
| Expt. id (Date / Name) | 2023-09-27 (b) | | | |
| Expt. type | Smart centrifugation | | | |
| Control plate count | 175 | 186 | 187 | 182.7 |
| Expt. plate count | 1 | 2 | 2 | 1.7 |
| Volume recovered (ml) | 2.7 | | | |
| Isolation efficiency (%) | 90.3 | | | |
| Concentration in blood (CFU/ml) | 22.1 | | | |
| Expt. id (Date / Name) | 2023-09-27 (c) | | | |
| Expt. type | Smart centrifugation | | | |
| Control plate count | 175 | 186 | 187 | 182.7 |
| Expt. plate count | 2 | 1 | 2 | 1.7 |
| Volume recovered (ml) | 2.6 | | | |
| Isolation efficiency (%) | 86.9 | | | |
| Concentration in blood (CFU/ml) | 22.1 | | | |
| Expt. id (Date / Name) | KP Chip Trail 6 | | | |
| Expt. type | Smart centrifugation + Microfluidic detection | | | |
| Control plate count | 4 | 4 | 3 | 3.7 |
| Expt. plate count | 13 | 13 | 16 | 14.0 |
| Volume recovered (ml) | 0.6 | | | |
| Isolation efficiency (%) | 132.7 | | | |

| | | | | |
|---|---|---|---|---|
| Concentration in blood (CFU/ml) | 7 | | | |
| Bacteria (t70) | 12 | | | |
| Bacteria (tend) | 14 | | | |
| relative isolation efficiency (%) (t70) | 76.2 | | | |
| Overall assay performance (%) (tend) | 88.9 | | | |
| Expt. id (Date / Name) | KP Chip Trail 3 | | | |
| Expt. type | Smart centrifugation + Microfluidic detection | | | |
| Control plate count | 30 | 28 | 12 | 23.3 |
| Expt. plate count | 133 | 134 | 85 | 117.3 |
| Volume recovered (ml) | 0.5 | | | |
| Isolation efficiency (%) | 99.4 | | | |
| Concentration in blood (CFU/ml) | 52 | | | |
| Bacteria (t70) | 65 | | | |
| Bacteria (tend) | 105 | | | |
| relative isolation efficiency (%) (t70) | 55.4 | | | |
| Overall assay performance (%) (tend) | 89.5 | | | |
| Expt. id (Date / Name) | 2024-02-27 (a) | | | |
| Expt. type | Smart centrifugation | | | |
| Control plate count | 206 | 210 | 222 | 212.7 |
| Expt. plate count | 4 | 2 | 2 | 2.7 |
| Volume recovered (ml) | 2.7 | | | |
| Isolation efficiency (%) | 120.4 | | | |
| Concentration in blood (CFU/ml) | 17.7 | | | |
| Expt. id (Date / Name) | 2024-02-27 (b) | | | |
| Expt. type | Smart centrifugation | | | |
| Control plate count | 206 | 210 | 222 | 212.7 |
| Expt. plate count | 1 | 1 | 3 | 1.7 |
| Volume recovered (ml) | 2.7 | | | |
| Isolation efficiency (%) | 75.2 | | | |
| Concentration in blood (CFU/ml) | 17.7 | | | |

| | | | | |
|---|---|---|---|---|
| Expt. id (Date / Name) | 2024-02-28 (a) | | | |
| Expt. type | Smart centrifugation | | | |
| Control plate count | 207 | 234 | 255 | 232.0 |
| Expt. plate count | 1 | 0 | 3 | 1.3 |
| Volume recovered (ml) | 2.5 | | | |
| Isolation efficiency (%) | 67 | | | |
| Concentration in blood (CFU/ml) | 14.7 | | | |
| Expt. id (Date / Name) | 2024-02-28 (b) | | | |
| Expt. type | Smart centrifugation | | | |
| Control plate count | 207 | 234 | 255 | 232.0 |
| Expt. plate count | 1 | 1 | 4 | 2.0 |
| Volume recovered (ml) | 2.5 | | | |
| Isolation efficiency (%) | 100.6 | | | |
| Concentration in blood (CFU/ml) | 14.7 | | | |

**Supplementary Table 11**: Raw data of plate counts of all experiments with *E. Faecalis*.

| | | | | Mean |
|---|---|---|---|---|
| Expt. id (Date / Name) | 2023-09-26 | | | |
| Expt. type | Smart centrifugation | | | |
| Control plate count | 7 | 10 | 22 | 13.0 |
| Expt. plate count | 7 | 9 | 3 | 6.3 |
| Volume recovered (ml) | 2.6 | | | |
| Isolation efficiency (%) | 46.4 | | | |
| Concentration in blood (CFU/ml) | 158 | | | |
| Expt. id (Date / Name) | 2024-02-29 (a) | | | |
| Expt. type | Smart centrifugation | | | |
| Control plate count | 20 | 26 | | 23 |
| Expt. plate count | 1 | 0 | 0 | 0.3 |
| Volume recovered (ml) | 2.5 | | | |
| Isolation efficiency (%) | 37.4 | | | |
| Concentration in blood (CFU/ml) | 9.9 | | | |
| Expt. id (Date / Name) | 2024-02-29 (b) | | | |
| Expt. type | Smart centrifugation | | | |
| Control plate count | 20 | 26 | | 23 |
| Expt. plate count | 1 | 0.0 | 0.0 | 0.3 |
| Volume recovered (ml) | 2.5 | | | |
| Isolation efficiency (%) | 37.4 | | | |
| Concentration in blood (CFU/ml) | 9.9 | | | |
| Expt. id (Date / Name) | 2023-09-27 (a) | | | |
| Expt. type | Smart centrifugation | | | |
| Control plate count | 94 | 96 | 96 | 95.3 |
| Expt. plate count | 99 | 89 | | 94.0 |
| Volume recovered (ml) | 2.7 | | | |
| Isolation efficiency (%) | 97.6 | | | |
| Concentration in blood (CFU/ml) | 1155.6 | | | |
| Expt. id (Date / Name) | 2023-09-27 (b) | | | |
| Expt. type | Smart centrifugation | | | |
| Control plate count | 94 | 96 | 96 | 95.3 |
| Expt. plate count | 94 | 86 | 78 | 86.0 |

| | | | | |
|---|---|---|---|---|
| Volume recovered (ml) | 2.7 | | | |
| Isolation efficiency (%) | 89.3 | | | |
| Concentration in blood (CFU/ml) | 1155.6 | | | |
| Expt. id (Date / Name) | 2023-09-28 (a) | | | |
| Expt. type | Smart centrifugation | | | |
| Control plate count | 101 | 106 | 106 | 104.3 |
| Expt. plate count | 7 | 9 | 9 | 8.3 |
| Volume recovered (ml) | 2.5 | | | |
| Isolation efficiency (%) | 86.5 | | | |
| Concentration in blood (CFU/ml) | 107 | | | |
| Expt. id (Date / Name) | 2023-09-28 (b) | | | |
| Expt. type | Smart centrifugation | | | |
| Control plate count | 101 | 106 | 106 | 104.3 |
| Expt. plate count | 9 | 9 | 11 | 9.7 |
| Volume recovered (ml) | 2.5 | | | |
| Isolation efficiency (%) | 100.4 | | | |
| Concentration in blood (CFU/ml) | 107.0 | | | |
| Expt. id (Date / Name) | 2023-09-28 (c) | | | |
| Expt. type | Smart centrifugation | | | |
| Control plate count | 101 | 106 | 106 | 104.3 |
| Expt. plate count | 1 | 1 | 0 | 0.7 |
| Volume recovered (ml) | 2.6 | | | |
| Isolation efficiency (%) | 60.9 | | | |
| Concentration in blood (CFU/ml) | 12.6 | | | |
| Expt. id (Date / Name) | 2023-09-28 (d) | | | |
| Expt. type | Smart centrifugation | | | |
| Control plate count | 101 | 106 | 106 | 104.3 |
| Expt. plate count | 1 | 0 | 0 | 0.3 |
| Volume recovered (ml) | 2.6 | | | |
| Isolation efficiency (%) | 30.5 | | | |
| Concentration in blood (CFU/ml) | 12.6 | | | |
| Expt. id (Date / Name) | 2023-09-28 (e) | | | |
| Expt. type | Smart centrifugation | | | |

| | | | | |
|---|---|---|---|---|
| Control plate count | 101 | 106 | 106 | 104.3 |
| Expt. plate count | 90 | 66 | 84 | 80.0 |
| Volume recovered (ml) | 2.5 | | | |
| Isolation efficiency (%) | 70.3 | | | |
| Concentration in blood (CFU/ml) | 1264.6 | | | |
| Expt. id (Date / Name) | 2024-03-01 | | | |
| Expt. type | Smart centrifugation | | | |
| Control plate count | 19 | 21 | 16 | 18.7 |
| Expt. plate count | 1 | 0 | 0 | 0.3 |
| Volume recovered (ml) | 2.6 | | | |
| Isolation efficiency (%) | 94.4 | | | |
| Concentration in blood (CFU/ml) | 4.1 | | | |
| Expt. id (Date / Name) | 2023-10-09 | | | |
| Expt. type | Smart centrifugation | | | |
| Control plate count | 15 | 23 | 19 | 19.0 |
| Expt. plate count | 1 | 2 | 2 | 1.7 |
| Volume recovered (ml) | 2.6 | | | |
| Isolation efficiency (%) | 83.6 | | | |
| Concentration in blood (CFU/ml) | 23 | | | |
| Expt. id (Date / Name) | 2024-02-29 (a) | | | |
| Expt. type | Smart centrifugation | | | |
| Control plate count | 20 | 26 | | 23.0 |
| Expt. plate count | 1 | 0 | 0 | 0.3 |
| Volume recovered (ml) | 2.6 | 2.5 | | |
| Isolation efficiency (%) | 37.4 | | | |
| Concentration in blood (CFU/ml) | 9.9 | | | |
| Expt. id (Date / Name) | 2024-02-29 (b) | | | |
| Expt. type | Smart centrifugation | | | |
| Control plate count | 20 | 26 | | 23 |
| Expt. plate count | 1 | 0 | 0 | 0.3 |
| Volume recovered (ml) | 2.6 | 2.5 | | |
| Isolation efficiency (%) | 37.4 | | | |
| Concentration in blood (CFU/ml) | 9.9 | | | |

| | | | | |
|---|---|---|---|---|
| Expt. id (Date / Name) | 2024-03-04 (a) | | | |
| Expt. type | Smart centrifugation | | | |
| Control plate count | 35 | 35 | 40 | 36.7 |
| Expt. plate count | 0 | 0 | 3 | 1.0 |
| Volume recovered (ml) | 2.6 | | | |
| Isolation efficiency (%) | 49.6 | | | |
| Concentration in blood (CFU/ml) | 23.3 | | | |
| Expt. id (Date / Name) | 2024-03-04 (b) | | | |
| Expt. type | Smart centrifugation | | | |
| Control plate count | 35 | 35 | 40 | 36.7 |
| Expt. plate count | 1 | 5 | 2 | 2.7 |
| Volume recovered (ml) | 2.6 | | | |
| Isolation efficiency (%) | 132.4 | | | |
| Concentration in blood (CFU/ml) | 23.3 | | | |
| Expt. id (Date / Name) | 2024-03-04 (c) | | | |
| Expt. type | Smart centrifugation | | | |
| Control plate count | 35 | 35 | 40 | 36.7 |
| Expt. plate count | 0 | 3 | 1 | 1.3 |
| Volume recovered (ml) | 2.6 | | | |
| Isolation efficiency (%) | 66.2 | | | |
| Concentration in blood (CFU/ml) | 23.3 | | | |
| Expt. id (Date / Name) | 2024-03-04 (d) | | | |
| Expt. type | Smart centrifugation | | | |
| Control plate count | 35 | 35 | 40 | 36.7 |
| Expt. plate count | 0 | 0 | 3 | 1.0 |
| Volume recovered (ml) | 2.6 | | | |
| Isolation efficiency (%) | 49.6 | | | |
| Concentration in blood (CFU/ml) | 23.3 | | | |
| Expt. id (Date / Name) | 2024-03-04 (e) | | | |
| Expt. type | Smart centrifugation | | | |
| Control plate count | 324 | 337 | | 330.5 |
| Expt. plate count | 11 | 13 | 8 | 10.7 |
| Volume recovered (ml) | 2.6 | | | |
| Isolation efficiency (%) | 58.7 | | | |

| | | | | |
|---|---|---|---|---|
| Concentration in blood (CFU/ml) | 209.8 | | | |
| Expt. id (Date / Name) | 2024-03-04 (f) | | | |
| Expt. type | Smart centrifugation | | | |
| Control plate count | 324 | 337 | | 330.5 |
| Expt. plate count | 11 | 8 | 9 | 9.3 |
| Volume recovered (ml) | 2.6 | | | |
| Isolation efficiency (%) | 51.4 | | | |
| Concentration in blood (CFU/ml) | 209.8 | | | |
| Expt. id (Date / Name) | 2024-03-04 (g) | | | |
| Expt. type | Smart centrifugation | | | |
| Control plate count | 324 | 337 | | 330.5 |
| Expt. plate count | 5 | 15 | 8 | 9.3 |
| Volume recovered (ml) | 2.6 | | | |
| Isolation efficiency (%) | 51.4 | | | |
| Concentration in blood (CFU/ml) | 209.8 | | | |
| Expt. id (Date / Name) | 2024-03-04 (h) | | | |
| Expt. type | Smart centrifugation | | | |
| Control plate count | 324 | 337 | | 330.5 |
| Expt. plate count | 15 | 13 | 7 | 11.7 |
| Volume recovered (ml) | 2.6 | | | |
| Isolation efficiency (%) | 64.2 | | | |
| Concentration in blood (CFU/ml) | 209.8 | | | |
| Expt. id (Date / Name) | 2024-03-04 (i) | | | |
| Expt. type | Smart centrifugation | | | |
| Control plate count | 324 | 337 | | 330.5 |
| Expt. plate count | 94 | 102 | 75 | 90.3 |
| Volume recovered (ml) | 2.6 | | | |
| Isolation efficiency (%) | 49.7 | | | |
| Concentration in blood (CFU/ml) | 2098.4 | | | |
| Expt. id (Date / Name) | 2024-03-04 (j) | | | |
| Expt. type | Smart centrifugation | | | |
| Control plate count | 324 | 337 | | 330.5 |
| Expt. plate count | 70 | 88 | 83 | 80.3 |

| | | | | |
|---|---|---|---|---|
| Volume recovered (ml) | 2.6 | | | |
| Isolation efficiency (%) | 44.2 | | | |
| Concentration in blood (CFU/ml) | 2098.4 | | | |
| Expt. id (Date / Name) | 2024-03-04 (k) | | | |
| Expt. type | Smart centrifugation | | | |
| Control plate count | 324 | 337 | | 330.5 |
| Expt. plate count | 94 | 113 | 90 | 99.0 |
| Volume recovered (ml) | 2.6 | | | |
| Isolation efficiency (%) | 54.5 | | | |
| Concentration in blood (CFU/ml) | 2098.4 | | | |
| Expt. id (Date / Name) | 2024-03-04 (l) | | | |
| Expt. type | Smart centrifugation | | | |
| Control plate count | 324 | 337 | | 330.5 |
| Expt. plate count | 74 | 76 | 100 | 83.3 |
| Volume recovered (ml) | 2.6 | | | |
| Isolation efficiency (%) | 45.9 | | | |
| Concentration in blood (CFU/ml) | 2098.4 | | | |
| Expt. id (Date / Name) | EF Chip trail 3 | | | |
| Expt. type | Smart centrifugation + Microfluidic detection | | | |
| Control plate count | 167 | 146 | 164 | 159.0 |
| Expt. plate count | 158 | 205 | 244 | 202.3 |
| Volume recovered (ml) | 0.6 | | | |
| Isolation efficiency (%) | 38.4 | | | |
| Concentration in blood (CFU/ml) | 1012 | | | |
| Bacteria (t70) | 167 | | | |
| Bacteria (tend) | 248 | | | |
| Overall assay performance (%) (tend) | 19.1 | | | |
| Expt. id (Date / Name) | EF Chip trail 4 | | | |
| Expt. type | Smart centrifugation + Microfluidic detection | | | |
| Control plate count | 23 | 51 | 52 | 42.0 |

| | | | | |
|---|---|---|---|---|
| Expt. plate count | 303 | 301 | 256 | 286.7 |
| Volume recovered (ml) | 0.6 | | | |
| Isolation efficiency (%) | 71.6 | | | |
| Concentration in blood (CFU/ml) | 143 | | | |
| Bacteria (t70) | 10 | | | |
| Bacteria (tend) | 10 | | | |
| Overall assay performance (%) (tend) | 4.3 | | | |
